# Differential roles of Na_V_1.2 and Na_V_1.6 in neocortical pyramidal cell excitability

**DOI:** 10.1101/2024.12.17.629038

**Authors:** Joshua D. Garcia, Chenyu Wang, Ryan P. D. Alexander, Emmie Banks, Timothy Fenton, Jean-Marc DeKeyser, Tatiana V. Abramova, Alfred L. George, Roy Ben-Shalom, David H. Hackos, Kevin J. Bender

**Affiliations:** Weill Institute for Neurosciences, University of California, San Francisco, San Francisco, CA, USA; Department of Neurology, University of California, San Francisco, San Francisco, CA, USA; Department of Neuroscience, Genentech, Inc., 1 DNA Way, South San Francisco, CA, USA; Department of Neurology, MIND Institute, University of California Davis School of Medicine, Sacramento, CA, 95817, USA; Department of Pharmacology, Northwestern University Feinberg School of Medicine, Chicago, IL USA

## Abstract

Mature neocortical pyramidal cells functionally express two sodium channel (Na_V_) isoforms: Na_V_1.2 and Na_V_1.6. These isoforms are differentially localized to pyramidal cell compartments, and as such are thought to contribute to different aspects of neuronal excitability. But determining their precise roles in pyramidal cell excitability has been hampered by a lack of tools that allow for selective, acute block of each isoform individually. Here, we leveraged aryl sulfonamide-based molecule (ASC) inhibitors of Na_V_ channels that exhibit state-dependent block of both Na_V_1.2 and Na_V_1.6, along with knock-in mice with changes in Na_V_1.2 or Na_V_1.6 structure that prevents ASC binding. This allowed for acute, potent, and reversible block of individual isoforms that permitted dissection of the unique contributions of Na_V_1.2 and Na_V_1.6 in pyramidal cell excitability. Remarkably, block of each isoform had contrasting—and in some situations, opposing—effects on neuronal action potential output, with Na_V_1.6 block decreasing and Na_V_1.2 block increasing output. Thus, Na_V_ isoforms have unique roles in regulating different aspects of pyramidal cell excitability, and our work may help guide development of therapeutics designed to temper hyperexcitability through selective Na_V_ isoform blockade.

## INTRODUCTION

Voltage-gated sodium channels (Na_V_) are critical for all aspects of neuronal excitability, from action potential (AP) initiation to axonal propagation, transmitter release, and dendritic excitability^1–6^. In mature neocortical pyramidal cells, electrogenesis and subsequent propagation of APs is systematically regulated with Na_V_ recruitment first initiated in the axon initial segment (AIS), with forward propagation along the axon and backpropagation into soma and dendrites^2,3,7^. This coordinated process is supported by membrane expression of the Na_V_ isoforms Na_V_1.6 and Na_V_1.2. Current models, based on immunostaining of channels and empirical measurements of excitability, suggest that Na_V_1.2 and Na_V_1.6 are differentially expressed across neurites. In the AIS, Na_V_1.6 predominates, with highest membrane density in the AP initiation region of the AIS most distal to the soma. Na_V_1.2, by contrast, has higher relative membrane density in the proximal AIS^2,8–12^. Along the axon, Na_V_1.6 appears enriched at nodes of Ranvier and terminals^13,14^. Somatodendritic densities are far lower, and current models suggest that the somatic and perisomatic membrane expresses equal levels of Na_V_1.6 and Na_V_1.2, whereas dendritic regions more distal to the soma are enriched exclusively with Na 1.2^4–6,15^. Thus, each channel isoform likely has a unique role in AP initiation and propagation simply based on their differential distribution in neuronal compartments.

To date, efforts to understand the unique contributions of different Na_V_ isoforms has relied largely on experiments in which channel genetic expression has been manipulated, either through constitutive or conditional knockout approaches. While these approaches have strong merit, interpretation is complicated by compensatory changes in other Na_V_ isoforms or other ion channel classes^4,5,16,17^. Pharmacological approaches, which are both acute and reversible, would be preferred, but identifying highly selective compounds that target particular Na_V_ isoforms is difficult, as *ScnXa* gene family isoforms have high degrees of amino acid sequence homology. Some compounds exhibit high potency, but differences in the half-maximal inhibitory concentration (IC_50_) for individual isoforms can be limited^18–21^. This is especially true for separation of Na_V_1.2 from Na 1.6, as these two channels have especially high sequence similarity^22,23^.

Aryl sulfonamide compounds (ASCs) constitute a unique class of Na_V_ inhibitors that potently bind activated Na_V_ isoforms, resulting in a stabilization of the inactivated state^24–28^. The binding pocket is shielded from ASCs when channels are closed due to the positioning of the S4 voltage sensing domain, but is bound rapidly by ASMs when exposed. Unbinding is promoted by strong hyperpolarization, but occurs much more slowly at physiological voltages (e.g., tau of 10^2^–10^3^ sec at −80 and −40 mV, respectively)^24^. In spiking neurons, this essentially imparts use-dependence to ASC block^24^, as the probability of channel block will increase in proportion with AP rates.

ASCs exhibit high potency for a subset of Na_V_ isoforms: Na_V_1.2, Na 1.6 and Na_V_1.7^24–27^. These isoforms share sequence homology at the binding pocket, each containing a tyrosine-tryptophan (YW) motif that helps stabilize ASC binding. By contrast, Na_V_1.1 and Na_V_1.3 harbor a serine-arginine (SR) sequence at the same site. Mutagenesis of Na_V_1.7 from YW to SR results in a 145-fold decrease in ASC binding without affecting channel biophysical properties^24^. Here, we engaged a similar strategy for Na_V_1.2 and Na_V_1.6, using knock-in mice with YW->SR substitutions in either or both channels. This allows for selective, potent ASC-mediated block of YW-containing channels while preserving the function of SR-containing channels. With these tools, we dissected the individual roles of Na_V_1.2 and Na_V_1.6 in neocortical pyramidal cell excitability. We found that Na_V_1.2 and Na_V_1.6 have unique—and at times conflicting—effects on overall AP excitability. Moreover, results suggest that ASCs can function as ‘on-demand’ pharmacological inhibitors that help normalize cellular excitability in seizure-like conditions. Together, this work highlights the importance of distinct Na_V_ isoforms localized to specific cellular domains and their respective contribution to AP properties to help facilitate activity across pyramidal neurons.

## RESULTS

### Aryl sulfonamides selectively bind and inhibit Na_V_ isoforms containing the YW motif

ASCs exhibit high affinity for an extracellular region within voltage sensing domain IV (VSD IV) of select Na_V_ channels with a conserved tyrosine-tryptophan (YW) motif^24^ (Fig. 1A, B). This YW motif found on Na_V_1.2, Na_V_1.6 and Na_V_1.7 allows for ASC stabilization following channel inactivation (Fig. 1B). By contrast, an SR motif present in Na_V_1.1 and Na 1.3 limits binding appreciably^24^. Thus, we hypothesized that converting either Na_V_1.2 or Na_V_1.6 channels from those that contain the YW motif to those that contain an SR motif would alter ASC binding to those channels significantly.

**Figure 1:**
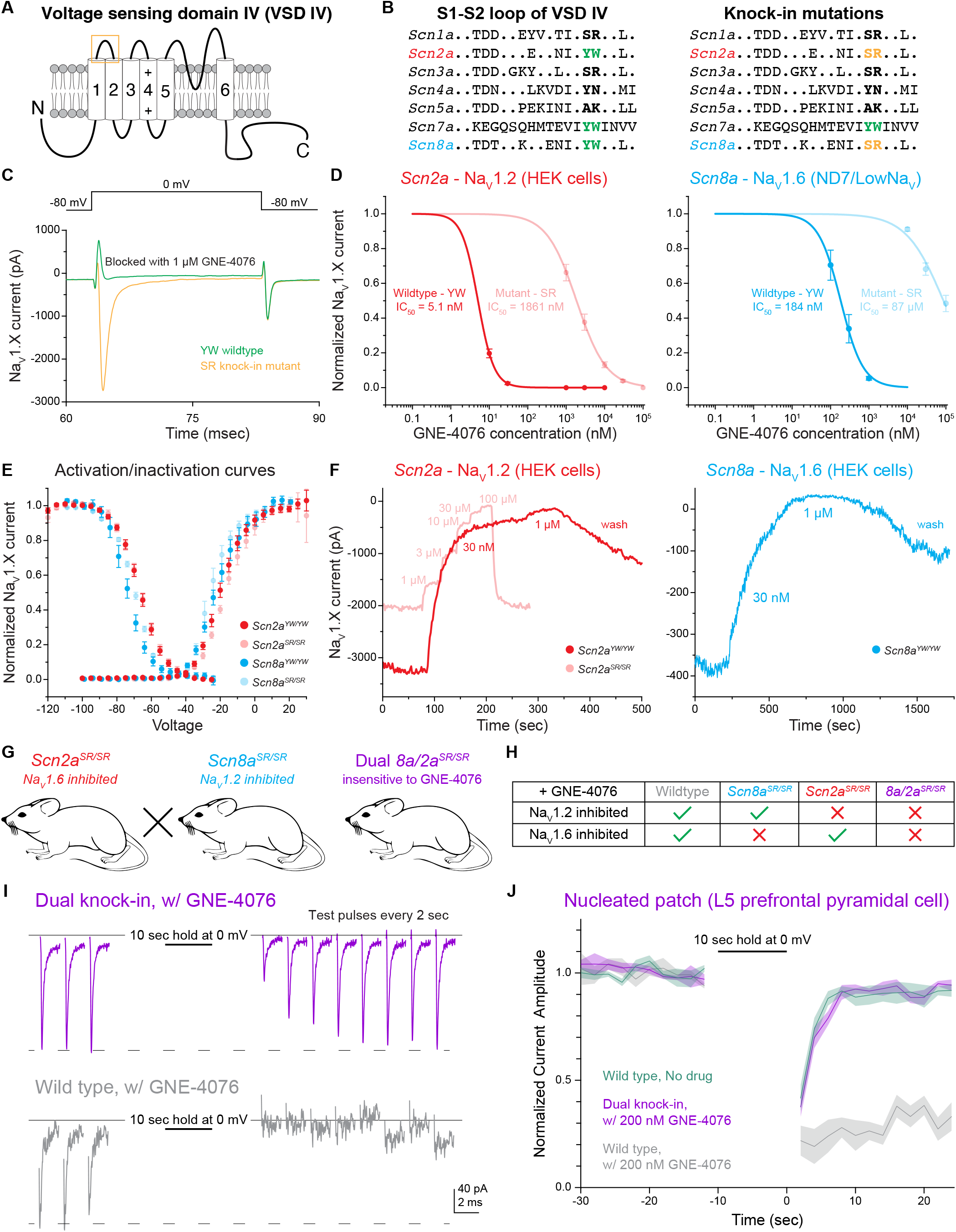
The YW motif on Na_V_1.2 and Na_V_1.6 increases activity-dependent GNE-4076 potency and subsequent channel inhibition. (A) Schematic depicting the fourth voltage sensing domain (VSD-IV) of Na_V_ isoforms. The six transmembrane spanning regions have high sequence homology amongst different Na_V_ isoforms, while linker regions display more sequence divergence. Orange box highlights extracellular S1-S2 loop where ASCs are stabilized by a tyrosine-tryptophan (YW) motif. (B) Amino acid sequence within the S1-S2 loop of various Na_V_ isoforms. *Scn2a* (Na_V_1.2) and *Scn8a* (Na_V_1.6) are the predominant channels expressed in mature, prefrontal pyramidal cells. Both isoforms share a conserved YW sequence that increases ASC potency. Knock-in mutations of *Scn2a* and *Scn8a* were generated by substituting the YW motif with a serine-arginine (SR) sequence present in *Scn1a* and *Scn3a*. (C) Example Na_V_ current traces (pA) of cells expressing either YW wildtype channels or SR knock-in mutant chimeras in the presence of 1 μM GNE-4076. To activate exogenously expressed Na_V_ channels, cells were held at −80 mV and stepped to 0 mV for 20 ms. GNE-4076 onboarding was performed by holding cells at 0 mV for 10 sec. (D) Dose response curves for exogenously expressed *Scn2a* (HEK cells) or *Scn8a* (ND7/LoNa_V_) in immortalized cell lines. IC_50_ was measured for both YW wildtype channels and SR knock-in mutant chimeras. YW->SR knock-in mutations reduced GNE-4076 potency by about 400-to 500-fold relative to wildtype channels. Circles represent normalized mean Na_V_ current amplitude ± SEM. (E) Activation and steady-state inactivation curves for both YW wildtype channels and SR knock-in mutant chimeras (*Scn2a*^*YW/YW*^, n=6; *Scn2a*^*SR/SR*^, n=7; *Scn8a*^*YW/YW*^, n=8; *Scn8a*^*SR/SR*^, n=6). *Scn2a* or *Scn8a* YW->SR mutations alter efficacy of GNE-4076 while having minor effects on biophysical properties of either isoform. Circles represent mean normalized Na_V_ current amplitude ± SEM. Unpaired t-test with Welch’s correction. No significance detected between wildtype and mutant channels for both V1/2 of activation or inactivation. (F) Example current amplitude response graphs for Na_V_1.2 (red) and Na_V_1.6 (blue) expressed in HEK cells. Cells were perfused with increasing concentrations of GNE-4076 throughout the recording. Individual current response recordings from HEK cells expressing *Scn2a* were robust (3.2 nA), and recordings were reproducible for both YW wildtype channels (red) and SR knock-in mutant chimeras (transparent red). Current responses from cells expressing *Scn8a* were variable with only a few cells exhibiting channel conductance (400 pA). In select cells expressing *Scn8a*, current amplitude (blue) also decreases substantially with 30 nM GNE-4076 and completely with 1 μM. (G) Transgenic mouse lines generated with the YW->SR knock-in mutation present on both *ScnXa* alleles. *Scn2a*^*SR/SR*^ mice were crossed with Scn8a^SR/SR^ mice to generate a dual *8a/2a*^*SR/SR*^ knock-in mouse. (H) Overview of the various transgenic (or wildtype) mouse lines used throughout this study. Application of 200 nM GNE-4076 selectively inhibits Na_V_ isoforms only containing the YW motif. (I) Nucleated patch experiments from prefrontal pyramidal cells performed in wildtype or dual 8a/2a^SR/SR^ knock-in cells in the presence of 200 nM GNE-4076. Baseline conductance was measured by depolarizing cells from −80 mV to −12 mV every 2 sec for 10 pulses. Na_V_ channels were inactivated by holding nucleated patch at −12 mV for 10 sec. Test pulses were again acquired during recovery similar to baseline pulses. (J) Summary graph of normalized current amplitude from nucleated patch experiments in (I). Baseline and recovery test pulses were acquired for at least 20 sec before and 25 sec after channel inactivation step. Solid line represents normalized mean Na_V_ current amplitude ± SEM. Graph also include wildtype, no drug control nucleated patch experiments (wildtype no drug, n=4; wildtype + GNE-4076, n=4; 8a/2a^SR/SR^ + GNE-4076, n=4).

To test for the effects of this motif substitution, we first examined currents generated by wild type and YW->SR mutated channels expressed in immortalized cell lines (Fig. 1C). HEK cells were primarily used for most experiments. But due to known low transfection efficiency of Na_V_1.6 in HEK cells, a subset of experiments were performed using an ND7/23 cell line engineered to lack most native Na_V_ conductance (ND7/LoNa)^35^. Both Na_V_1.2 and Na_V_1.6 wildtype channels expressed in cell lines were inhibited markedly by 1 μM GNE-4076 (Fig. 1C). By contrast, SR knock-in mutants continued to flux sodium in the presence of 1 μM GNE-4076. Dose response curves revealed an IC_50_ of 5.1 nM for wildtype Na_V_1.2 that increases 365-fold to 1861 nM for mutant SR channels (Fig. 1D). For Na_V_1.6 channels, GNE-4076 IC_50_ increased 475-fold from 184 nM to 87 μM with the SR mutant (Fig. 1D). To assess whether SR mutations affect channel gating properties in the absence of GNE-4076 binding, steady state activation and inactivation curves were assessed for wildtype or mutant channels (Fig. 1E). Consistent with prior work studying a similar mutation in Na 1.7^24^, the YW to SR mutation had no effect on slope factor for both voltage-dependent activation or inactivation for either Na_V_1.2 or Na_V_1.6 (Table 2); however, both voltage-dependent activation and inactivation hyperpolarized by approximately 2 mV for Na_V_1.6, but not Na_V_1.2.

While GNE-4076 binds potently to either Na_V_1.2 (Fig. 1D, F) or Na_V_ 1.7 expressed in HEK cells^28^, its affinity was lower for Na 1.6 expressed in ND7/LoNa_V_ cells. To test if this was due to reductions in affinity imposed by the ND7/23 cell line, we examined current in the few HEK cells in which Na_V_1.6 current could be measured (Fig. 1F). While peak currents were small (400 pA vs 3.2 nA for Na_V_1.2 transfected using identical protocols), we found that 30 nM GNE-4076 exhibited strong block of Na_V_1.6 (Fig. 1F), but due to limitations in expression efficiency in HEK cells, we were unable to collect full dose-response curves to determine IC_50_.

Results in heterologous expression systems indicate that YW −>SR substitution reduces GNE-4076 binding affinity markedly. Given these results, we constructed transgenic mice containing the YW->SR mutation. Two distinct lines, termed *Scn8a*^*SR/SR*^ and *Scn2a*^*SR/SR*^, were generated by mutating both alleles of each sodium channel gene (Fig. 1G). SR/SR mutations were validated by PCR (see Methods) and mice were bred to be homozygous for the mutations. Comparisons were then made between wild type mice, mice with YW->SR mutations into Na_V_1.2 or Na_V_1.6 alone (*2a*^*SR/SR*^ or *8a*^*SR/SR*^, respectively), or YW->SR mutations in both Na 1.6 and Na_V_ 1.2 (dual *8a/2a*^*SR/SR*^). In principle, one could determine a concentration of GNE-4076 in which native channels are blocked potently, sparing SR variant channels (Fig. 1H). Thus, these mice should enable selective, acute manipulations of Na_V_1.2 and Na_V_1.6-dependent aspects of neuronal excitability using a chemical genetics approach.

Given the disparate binding results in heterologous cell lines, we first tested the effects of a dose that should potently block WT channels but not affect SR variant channels to verify that native channels behave similarly to those expressed in HEK cells. Neocortical pyramidal cell somata are thought to express similar levels of Na_V_1.2 and Na_V_1.6 on their membranes^4,5,8^. Thus, excised patches from this region can provide insight into block of both isoforms. Acute coronal slices containing mPFC were prepared from WT or *8a/2a*^*SR/SR*^ mice and nucleated patches were excised from layer 5 pyramidal cells. After establishing a stable baseline of Na_V_-mediated current evoked from a holding voltage of −80 mV, a value comparable to the resting membrane potential studied in later current-clamp experiments, cells were pulsed to −12 mV for 10 sec to achieve near-complete block of NaV isoforms, then returned to −80 mV (Fig. 1I). In untreated WT conditions, currents recovered to near-baseline levels within 5 sec (Fig. 1J). A similar quick recovery was observed in cells from *8a/2a*^*SR/ SR*^ mice in the presence of 200 nM GNE-4076. By contrast, WT neurons exposed to the same dosing were inhibited markedly, with no appreciable recovery within 25 sec (Fig. 1I, J). Taken together, these results show that the YW to SR mutation reduces ASC binding and inhibition of Na_V_ chimeras both in immortalized cell lines and pyramidal neurons.

We also asked whether neuronal AP properties were affected by these mutations in *8a/2a*^*SR/SR*^ mice by assessing both threshold and peak dV/dt in the absence of GNE-4076 (Fig. S1). Compared to wildtype cells, there were no detectable changes to peak dV/dt in *8a/2a*^*SR/SR*^ neurons (Fig. S2B). However, AP threshold was hyperpolarized in *8a/2a*^*SR/SR*^ neurons, likely due to small changes in voltage-dependence of activation observed for heterologously expressed Na_V_1.6 (Fig. S2A, Table 2). Application with GNE-4076 in either wildtype or *8a/2a*^*SR/SR*^ neurons had no further effect on threshold or peak dV/dt compared to controls recording in the absence of GNE-4076, suggesting minimal drug binding occurs without marked depolarization or spiking activity (Fig. S1, Table 3).

### Differential roles for Na_V_1.6 and Na_V_1.2 in AP initiation and somatic excitability

Na_V_1.2 and Na_V_1.6 are differentially distributed in neuronal arbors and are thought to contribute to different aspects of AP initiation and propagation. In mature neocortical pyramidal cells, Na_V_1.6 is enriched in the distal AIS, the site where APs initiate^4,8,16,44^. Following initiation, APs forward propagate along the axon via Na_V_1.6-enriched nodes of Ranvier, but also backpropagate into the soma through a region enriched with a mix of Na_V_ 1.2 and Na_V_ 1.6^8^. Several studies have sought to identify specific roles for each isoform by conditional deletion of either isoform^4,16^. Unfortunately, in such conditions, the residual isoform compensates for loss to some degree, making interpretation of individual isoform roles difficult. We therefore leveraged ASC-based block to study acute, differential inhibition of Na_V_ isoforms to better understand the individual roles of Na_V_1.2 and Na_V_1.6 in AP excitability.

Changes to AP waveform were visualized with phase plane plots, which plot membrane voltage vs. membrane voltage velocity (Fig. 2B and S3A). Within phase plots, spike threshold appears as a sudden deviation from rest and is defined as the voltage at which voltage velocity (dV/dt) first exceeds 15 V/s (Fig. S3A). Following spike threshold, a neuron’s voltage transits through two components of Na_V_ recruitment, first in the AIS and then in the soma^7,45^. These result in characteristic “humps” in the depolarizing aspect of the phase plot^2,11^. Thus, these different components of the phase plot can aid one’s understanding of effects on specific components of Na_V_-mediated excitability.

**Figure 2:**
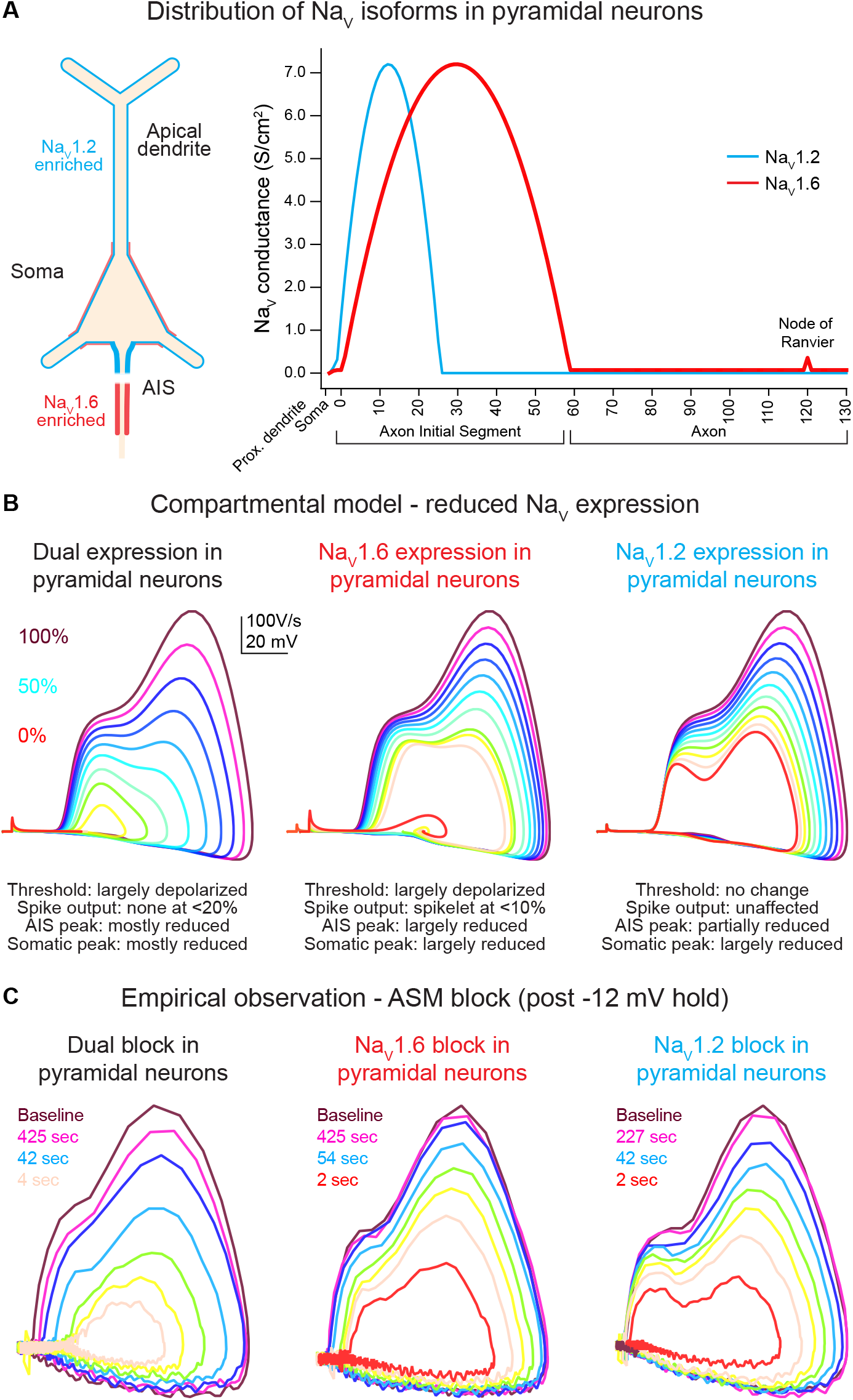
Global *Scn2a, Scn8a* or dual loss in compartmental models distinctly impacts key AP properties. (A) Na_V_1.6 and Na_V_1.2 are equally expressed in soma and proximal dendrites. Expression pattern is more distinct in other regions with Na_V_1.6 enriched in the distal AIS, axon and nodes of Ranvier, whereas Na_V_1.2 is found exclusively in the proximal AIS and distal dendrites. (B) Compartmental model representing changes to phase plots when Na_V_ isoform expression is reduced from 100% (warmer colors) to 0% (in 10% increments) based on known localization across distinct neuronal localities. Lower Na_V_1.6 expression depolarized spike threshold and decreased both AIS and somatic AP velocity (dV/dt). Reduced Na_V_1.2 expression largely impacts backpropagation and somatic AP velocity. (C) Empirical observations of phase plot following near complete channel block with ASCs. Darker trace represents phase plot taken at baseline prior to −12 mV hold for 30 sec (see Fig. 3A). Colored traces represent recovery phase plots at different times post −12 mV hold for 30 sec with warmer colors depicting more time lapsed and increased channel recovery.

To provide *a priori* predictions of potential ASC-based effects on excitability, we constructed a compartmental model in which each channel’s density could be modulated. Channels were distributed based on predictions from empirical anatomical and physiological studies^4,6,8,16,46^. Na_V_1.6 was enriched in the distal AIS, Na_V_1.2 was enriched in the proximal AIS, both channels were expressed at equal levels in the soma and the first 20 microns of dendrite closest to the soma, and Na_V_1.2 was expressed exclusively in all other dendrites (Fig. 2A). Within this model, each channel’s density was modulated in 10% increments, from 100 to 0% (Fig. 2B).

In WT conditions (100% density of both channels), models generated APs with a threshold and AP kinetics comparable to empirical baseline observations (Fig. 2B, C). Within this model, reducing Na_V_1.2 or Na_V_1.6 density had clear, dissociable effects. Progressive reduction of Na_V_1.6 produced a progressive depolarization of AP threshold and a corresponding decrement in dV/dt throughout the entire rising phase of the AP. Na_V_1.2 reduction, by contrast, had no effect on AP threshold or the initial velocity of the AP initiated in the distal AIS. Instead, components of the AP related to backpropagation and recruitment of somatic Na_V_ channels were impaired only, with a decrement in peak dV/dt and peak AP membrane potential. To validate the accuracy of the parameters used in our model, we also performed sensitivity analysis (Fig. S2). Here, we adjusted the crossover point of Na_V_1.2 and Na_V_1.6 within the AIS as well as varied the density ratio of both channels (Fig. S2A, B). Increasing Na_V_1.6 ratio to 100% consistently hyperpolarized AP threshold across all AIS crossover positions (Fig. S2B, C). However, dV/dt was highly variable and random when either the AIS crossover point is shifted or Na_V_ ratios are altered.

While these models provide clues to the differential roles of Na_V_1.2 and Na_V_1.6, we needed to design an experimental strategy that increased the overall degree of Na_V_ blockade using ASCs in mice where either Na_V_1.6 or Na_V_1.2 was mutated to be insensitive to 200 nM GNE-4076 binding (Fig. 3C, 1G). Since it is nearly impossible to achieve full block with ASCs under physiological firing, we developed a hybrid current- and voltage-clamp experiment to study effects of more complete isoform blockade (Fig. 3A). In this protocol, neurons were held to −80 mV in current clamp with constant bias current (if necessary) and baseline APs were elicited with brief somatic current injection (300 ms, amplitude adjusted to evoke ~4-5 APs). We then promoted Na_V_ activation and inactivation by voltage-clamping neurons to −12 mV for 30 sec (Fig. 3A). Following voltage-clamp, neurons were returned to current-clamp with the same bias current as used in baseline conditions. We then evoked APs, first at an interstimulus interval of 2 sec, then at successively increasing intervals of 5 to 60 sec, allowing for somewhat consistent sampling on a log-base timescale that aligns well with both channel recovery from inactivation and recovery from GNE-4076 block (Fig. 3D, F). For clarity and simplicity, we will describe studies based on which channel is inhibited rather than which channel was rendered insensitive to GNE-4076. For example, a study in the *Scn2a*^*SR/SR*^*/Scn8a*^*+/+*^ animal is a case where Na_V_ 1.6 can be blocked.

**Figure 3:**
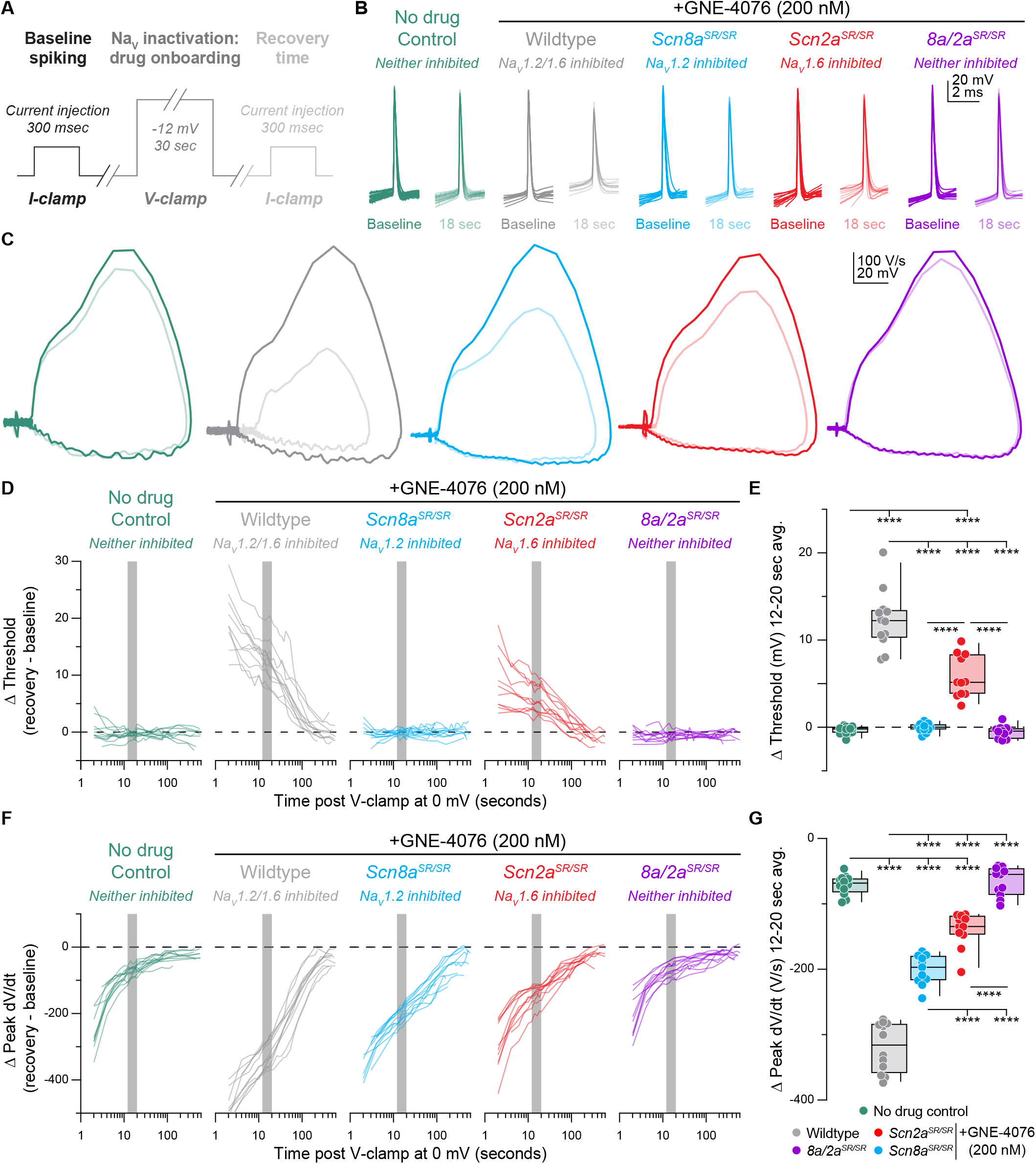
Recovery of AP firing properties is greatly diminished following dual Na_V_1.6 and Na_V_1.2 inhibition compared to selective block of individual channels. (A) Protocol used to characterize recovery of AP firing properties. Baseline spiking is determined by injecting current for 300 ms to elicit 5-6 APs. To promote Na_V_ inactivation and maximal GNE-4076 onboarding, neurons are held at −12 mV in voltage-clamp for 30 sec. Recovery of AP firing is evaluated by injecting same current stimulus defined during baseline spiking with an inter-stimulus interval starting at 2 sec, followed by 5,15, 30 and 60 sec. (B) Overlaid waveform of 1st AP at baseline or 18 sec post GNE-4076 onboarding for all conditions (wildtype no drug, n=12; wildtype + GNE-4076, n=11; *8a/2a*^*SR/SR*^ + GNE-4076, n=12; *Scn2a*^*SR/SR*^ + GNE-4076, n=12; *Scn8a*^*SR/SR*.^ + GNE-4076, n=12). (C) Overlaid phase plane of AP traces at baseline (100% transparency) or 18 sec post GNE-4076 onboarding (20% transparency) for each condition in (B). Plots represent the AP velocity by taking the first derivative (dV/dt, y-axis) versus the membrane potential (mV, x-axis). Colors are matched to conditions represented in (B). (D) Recovery of AP threshold (V_m_) represented as a delta value for individual cells plotted against time post GNE-4076 onboarding (log-scale). For Δ V_m_, baseline value is subtracted from individual timepoints throughout the recovery phase (Δ mV = recovery timepoint – baseline). Colors are matched to conditions represented in (B). Gray shaded bar represents recovery between 12-20 sec. (E)Summary data for Δ V_m_ at 12-20 sec post GNE-4076 onboarding (time period represented as gray bar in (D)). Box plots show median and 90% tails. Circles represent individual cells. One-way ANOVA, Holm-Šídák multiple comparisons test. ****p<0.0001.

Empirical data at various timepoints post −12 mV hold reveals that recovery from ASC-dependent block increases channel availability from approximate 0-10% to about 90-95% of the baseline spike when more time has elapsed (Fig. 2C). This recovery in available channels mimics changes observed to phase plots in compartmental models based on overall channel expression (Fig. 2B).

We then focused on our analysis on recovery of threshed or peak dV/dt across all genotypes (Fig. 3). In untreated WT cells, AP threshold following voltage-clamp recovered to baseline values immediately (Fig. 3C, D). By contrast, peak somatic dV/dt recovered more slowly (Fig. 3C, F), likely reflecting recovery from slow inactivation of channels in the perisomatic region^47–49^. Peak dV/dt recovered to within 87.12 ± 0.86% by 12-20 sec of voltage-clamp offset (Fig. 3F, G). Subsequent recovery of the residual peak dV/dt was slower, taking another 15-30 sec to recover the next 5% of peak dV/dt (Fig. 3F). Identical results were observed in *8a/2a*^*SR/SR*^ cells treated with 200 nM GNE-4076, suggesting that any effect observed in single knock-in recordings will be due to block of GNE-4076 sensitive channels (Fig. 3B-G). We therefore focused on the 12-20 sec after voltage-clamp offset for subsequent analysis, as it is a period in which most channel-intrinsic recovery has occurred, but also a period in which we would still expect significant block from GNE-4076.

When individual channel isoforms were blocked more completely with voltage steps to −12 mV, dramatic changes in AP threshold and peak dV/dt were observed. When Na_V_1.6 was blocked, threshold depolarized by 5.8 ± 0.7 mV (Fig. 3E). When Na_V_1.2 was blocked instead, threshold was unaffected (Δ Vm: −0.1 ± 0.2). Thus, AP threshold and AP initiation appears to be initiated in an Na_V_1.6-rich region in control conditions; but when Na_V_1.6 is inhibited, APs can occur at more depolarized potentials, likely mediated predominately by Na_V_1.2.

To examine these relative contributions further, we analyzed the rising phase of APs more closely (Fig. S3A). As described above, the first component of the rising phase of the AP reflects recruitment of Na_V_ channels localized to the AIS (Fig. S3A, middle). Based on changes in voltage acceleration (second derivative) within this period, the AIS component can be further divided into initiation and AIS backpropagation components^8,37^ (Fig. S3A, right). These periods were divided based on a timepoint during the AIS phase where voltage acceleration first peaked (AIS inflection point) or afterwards decreased, creating a trough in an acceleration vs. time graph (AIS max; Fig. S3B). The dV/dt at the AIS inflection point was affected only by Na_V_1.6 block (Fig. S3C), whereas the dV/dt at the trough was affected by block of either isoform (Fig. S3D, E). This reinforces the model where APs are initiated via the Na_V_1.6-enriched distal AIS, and where the depolarization produced by these distally localized channels recruits Na_V_1.2 in the more proximal AIS. Absolute values are also reported for threshold, peak dV/dt and AIS max for all recorded cells in recovery experiments (Fig. S4).

### Acute block of Na_V_1.2 increases pyramidal cell AP output

Previously, we showed that conditional knockout of *Scn2a* increased AP output^4^. We hypothesized that this was due to the lack of Na_V_ 1.2 in dendrites. When they are absent from this compartment, the dendrite does not depolarize as effectively, leading to a corresponding decrease in the recruitment of dendrite-localized potassium channels (K_V_). Consequently, neurons repolarize less between APs, making it easier to evoke the next AP in the Na_V_1.6-enriched AIS.

Though this hypothesis was supported by compartmental modeling demonstrating that acute block of Na_V_1.2 mirrored empirical observations, we could not eliminate the possibility that some form of cellular compensation occurred in the weeks between conditional knockout induction and acute slice experiments^50^. Therefore, we leveraged GNE-4076 to test the effects of acute Na_V_1.2 block (Fig. 4A), eliminating the possibility of genetic or post-translational compensation. We further compared this manipulation to block of Na_V_1.6 or both isoforms by examining overall AP output from protocols described in Fig. 3A.

**Figure 4:**
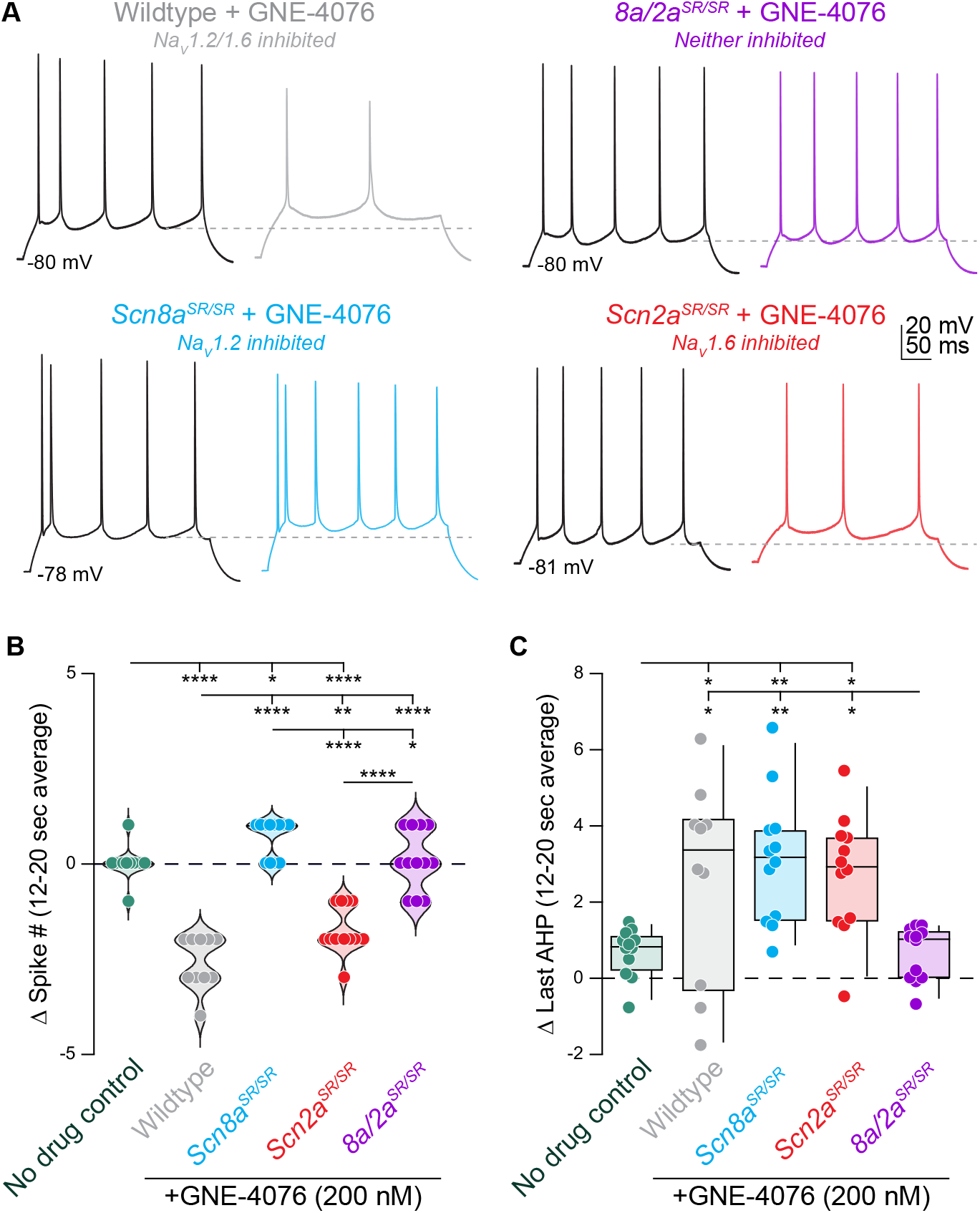
Acute inhibition of Na_V_1.2 increases AP excitability. (A) AP train over 300 ms at baseline (black) or 18 sec post GNE-4076 onboarding (color) for all conditions (wildtype + GNE-4076, n=10; *8a/2a*^*SR/SR*^ + GNE-4076, n=12; *Scn2a*^*SR/ SR*^ + GNE-4076, n=12; *Scn8a*^*SR/SR*.^ + GNE-4076, n=12). Dashed line represents V_m_ of last after-hyperpolarization (AHP). (B) Summary data for Δ spike number at 12-20 sec post GNE-4076 onboarding. One-way ANOVA, Holm-Šídák multiple comparisons test. *p<0.05, **p<0.01, ****p<0.0001. (C) Summary data for Δ last AHP at 12-20 sec post GNE-4076 onboarding. One-way ANOVA, Holm-Šídák multiple comparisons test. *p<0.05, **p<0.01. **(F)** Recovery of AP peak velocity (dV/dt) represented as a delta value for individual cells plotted against time post GNE-4076 onboarding (log-scale). For Δ dV/dt, baseline value is subtracted from individual timepoints throughout the recovery phase (Δ V/s= recovery timepoint – baseline). Colors are matched to conditions represented in (B). Gray shaded bar represents recovery between 12-20 sec. (G) Summary data for Δ dV/dt at 12-20 sec post GNE-4076 onboarding (time period represented as gray bar in (F)). Box plots show median and 90% tails. Circles represent individual cells. One-way ANOVA, Holm-Šídák multiple comparisons test. ****p<0.0001.

Remarkably, acute Na_V_1.2 block mirrored conditional knockout, and was the only condition in which AP output increased (Fig. 4B) and was associated with a depolarization in afterhyperpolarization voltage (Fig. 4C). In contrast, Na_V_1.6 block decreased AP output, as did block of both channels (Fig. 4B). This indicates that blocking Na_V_1.2 alone can increase AP output, independent of compensatory changes to other channels that may occur with genetic manipulations.

### Activity-dependent effects of ASCs on Na_**V**_ channels and subsequent inhibition alters AP properties and firing rate

During physiological activity, non-selective pharmacological inactivated state^24–27^. In contrast to data presented above where we observe near complete block immediately following prolonged depolarization (Fig. S4, absolute values), neurons experiencing physiological levels of activity likely never reach high percentages of channel blockade. Thus, in normal spiking neurons, ASCs would therefore be predicted to exhibit use-dependence, progressively blocking channels in proportion to a neuron’s activity rate.

To test this concept, we generated prolonged periods of activity in current-clamp by injecting 300 pA into the somatic pipette for 10 sec (Fig. 5A). Drug-naïve WT neurons responded to somatic current injection with inhibition of Na_V_ and AP firing channels in neurons greatly reduces cellular excitability repetitive spiking at a rate of 14.7 ± 0.5 Hz (n = 12). Neurons fired at a steady state after the first second, with stable AP threshold and instantaneous AP ^51,52^. ASCs are unique in that they require prolonged bursts of activity or even neuronal hyperexcitability to stabilize more channels in the frequency (Fig. 5C). Spiking characteristics were identical in *8a/2a*^*SR/SR*^ cells treated with GNE-4076, indicating that, at 200 nM, neurons are insensitive to GNE-4076 when the YW->SR motif substitution is present (Fig. 5C, D). However, in WT neurons treated with GNE-4076, threshold continued to depolarize after the first second, with corresponding decrements in instantaneous frequency and peak AP dV/dt throughout the stimulus (Fig. 5C, D). This demonstrates that GNE-4076 suppresses firing of neurons with a baseline firing rate of ~15 Hz (Table 4).

**Figure 5:**
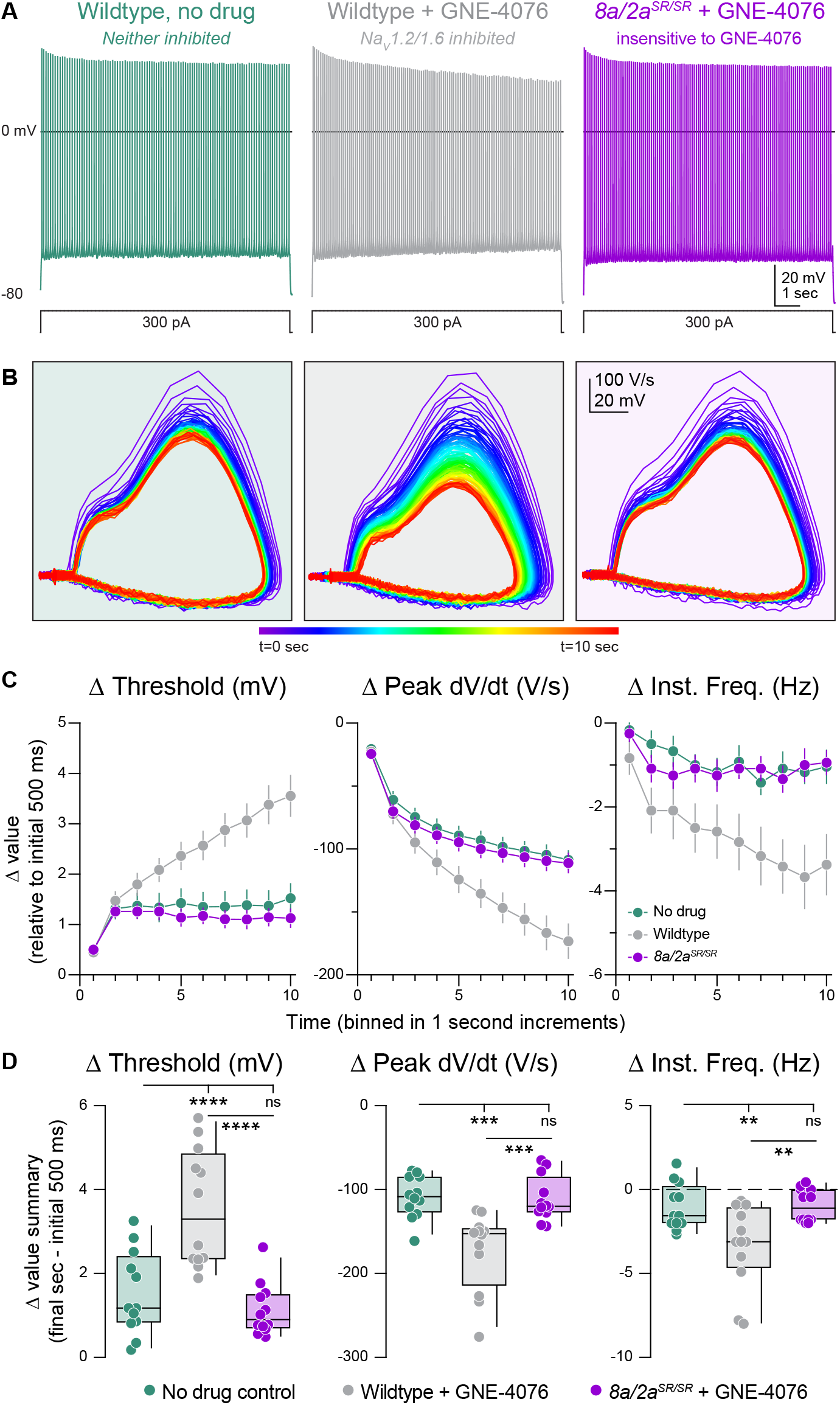
Activity-dependent onboarding of GNE-4076 to Na_V_ channels alters action potential firing properties in layer 5b, thick-tufted excitatory neurons. (A) Representative AP firing response to 300 pA current injection for 10 sec in wildtype or 8a/2a^SR/SR^ cells with or without 200 nM GNE-4076. (B) Phase plane of AP traces shown in (A). Plots represent AP velocity by taking the first derivative (dV/dt, y-axis) versus the membrane potential (mV, x-axis). To represent changes with phase plane relative to time, a rainbow color spectrum is used with warmer colors representing more time lapsed. (C) Delta threshold (Δ mV), delta peak dV/dt (Δ V/s) and delta instantaneous firing frequency (Δ Hz) binned in 1 sec increments normalized to the initial 500 ms of current injection (binned time – initial 500 ms). Circles represent mean Δ value ± SEM. Two-way ANOVA, Holm-Šídák multiple comparisons test. (D) Summary data for the final sec in (C). Delta values are normalized to the initial 500 ms of the stimulus (binned time – initial 500 ms). Box plots show median and 90% tails. Circles represent individual cells (wildtype no drug, n=12; wildtype + GNE-4076, n=12; 8a/2a^SR/SR^ + GNE-4076, n=12). One-way ANOVA, Holm-Šídák multiple comparisons test. **p<0.01, ***p<0.001, ****p<0.0001.

We then repeated experiments described above—where APs were generated with somatic depolarization over 10 sec (Fig. 6). In cases where Na_V_1.6 alone could be inhibited, AP threshold depolarized in response to GNE-4076 as much as in WT cases (Fig. 6C, D). Furthermore, AP frequency and peak somatic AP dV/dt was reduced to levels observed in WT cells (Fig. 6C, D). In cases where Na_V_1.2 alone could be inhibited, changes in AP threshold and instantaneous frequency were no different than *8a/2a*^*SR/SR*^ conditions. Indeed, the only change in AP properties was a decrease in peak AP dV/dt (Fig. 6D). This suggests that AP threshold and instantaneous frequency are more sensitive to Na_V_1.6 antagonism, whereas peak somatic dV/dt can be affected by antagonism of either isoform. Absolute values are also reported for threshold, peak dV/dt and instantaneous frequency for all recorded cells in prolonged activity burst experiments (Fig. S5).

**Figure 6:**
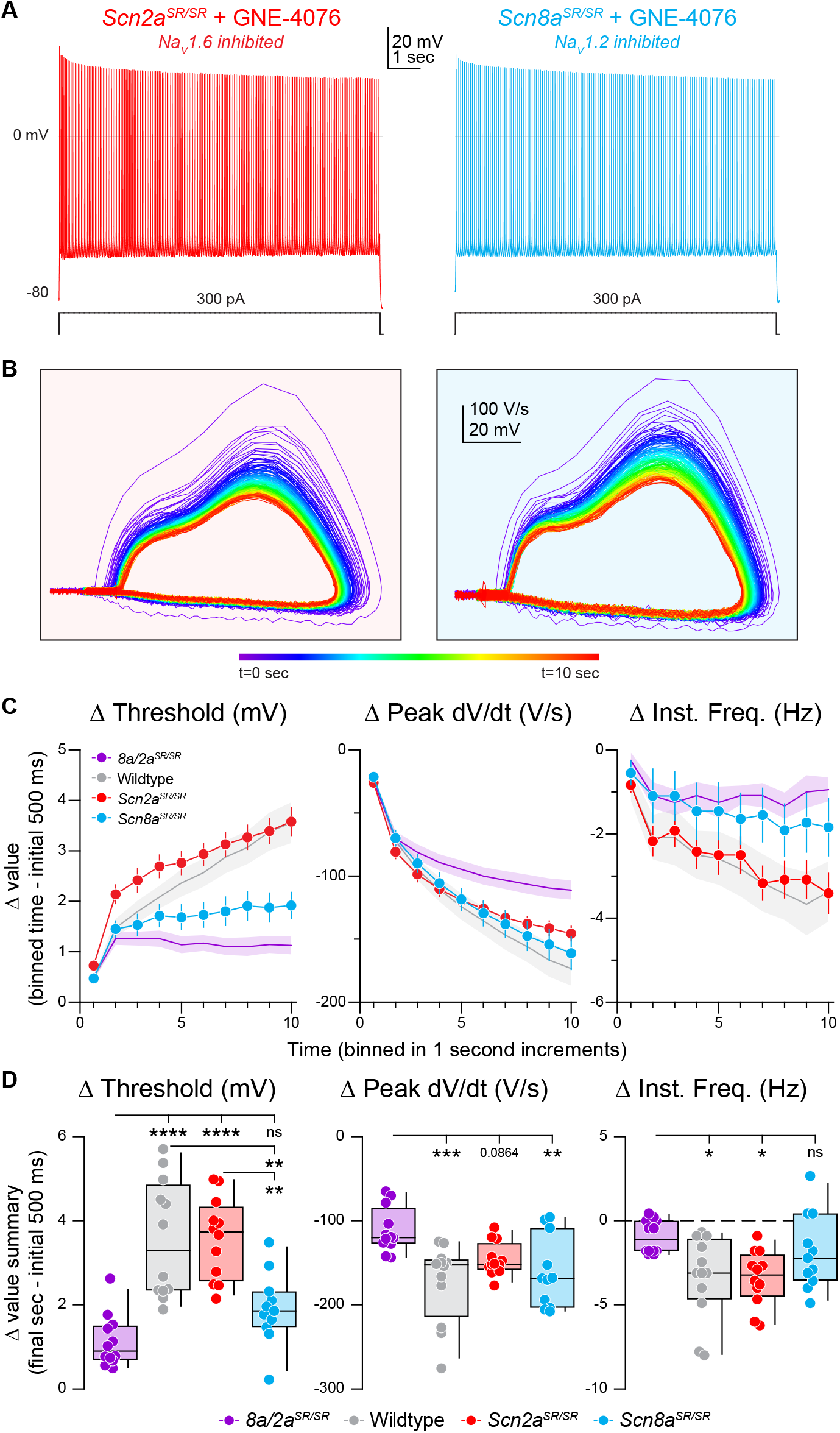
Selective inhibition of Na_V_1.6 depolarizes AP threshold markedly while blocking both Na_V_1.6 and Na_V_1.2 reduces AP velocity. (A) Representative AP firing response to 300 pA current injection for 10 sec in *Scn2a*^*SR/SR*^ or *Scn8a*^*SR/SR*^ cells with 200 nM GNE-4076 to selectively inhibit Na_V_1.6 or NaV1.2, respectively. (B) Phase plane of AP traces shown in (A). Plots represent AP velocity by taking the first derivative (dV/dt, y-axis) versus the membrane potential (mV, x-axis). To represent changes with phase plane relative to time, a rainbow color spectrum is used with warmer colors representing more time lapsed. (C) Delta threshold (Δ mV), delta peak dV/dt (Δ V/s) and delta instantaneous firing frequency (Δ Hz) binned in 1 sec increments normalized to the initial 500 ms of current injection (binned time – initial 500 ms). Circles represent mean Δ value ± SEM. Average Δ value ± SEM for *8a/2a*^*SR/ SR*^ + GNE-4076 and wildtype + GNE-4076 from Fig. 2D are represented. (D) Summary data for the final sec in (C). Delta values are normalized to the initial 500 ms of the stimulus (binned time – initial 500 ms). Box plots show median and 90% tails. Circles represent individual cells (*8a/2a*^*SR/SR*^ + GNE-4076, n=12; wildtype + GNE-4076, n=12; *Scn2a*^*SR/SR*^ + GNE-4076, n=10; *Scn8a*^*SR/SR*^ + GNE-4076, n=12). One-way ANOVA, Holm-Šídák multiple comparisons test. *p<0.05, **p<0.01, ***p<0.001, ****p<0.0001.

### Leveraging use-dependent, isoform-selective Na_**V**_ pharmacology as anticonvulsants

Epilepsy is often associated with aberrant, excessive excitability in neocortical networks, whether from direct hyperexcitability in pyramidal cells^53,54^ or disinhibition of pyramidal cells via alterations in inhibitory networks^37,55^. Several use-dependent Na antagonists with structures similar to GNE-4076 are being developed, exhibiting differential selectivity for Na_V_ 1.2 and Na_V_ 1.6^25^. Use-dependence may have advantages as antiepileptics. In theory, they would have minimal effect on neurotypical network activity unless such networks were hyperactive enough to promote drug binding. This may occur preferentially in seizure states. But given results above demonstrating that block of Na_V_1.2 and Na_V_1.6 can have different effects on overall AP output, it is critical to determine how block of either channel affects overall activity in seizure-like conditions.

To test this, we mimicked synaptic input by repeatedly injecting a 60 sec-long somatic current composed of Poisson-distributed EPSC and IPSC-like waveforms designed to elicit ~10 Hz spiking in baseline conditions (Fig. 7A). Following this baseline, the PSC-like train was repeated, imposed upon a 400 pA standing current injection to increase spiking to ~30 Hz (Fig. 7C), mimicking spike rates commonly observed during cortical seizure^56,57^. After this seizure-like event, cells were returned to baseline voltages and recovery was assessed with 4 more repetitions of the 60 sec PSC stimulus (Fig. 7A-C).

**Figure 7:**
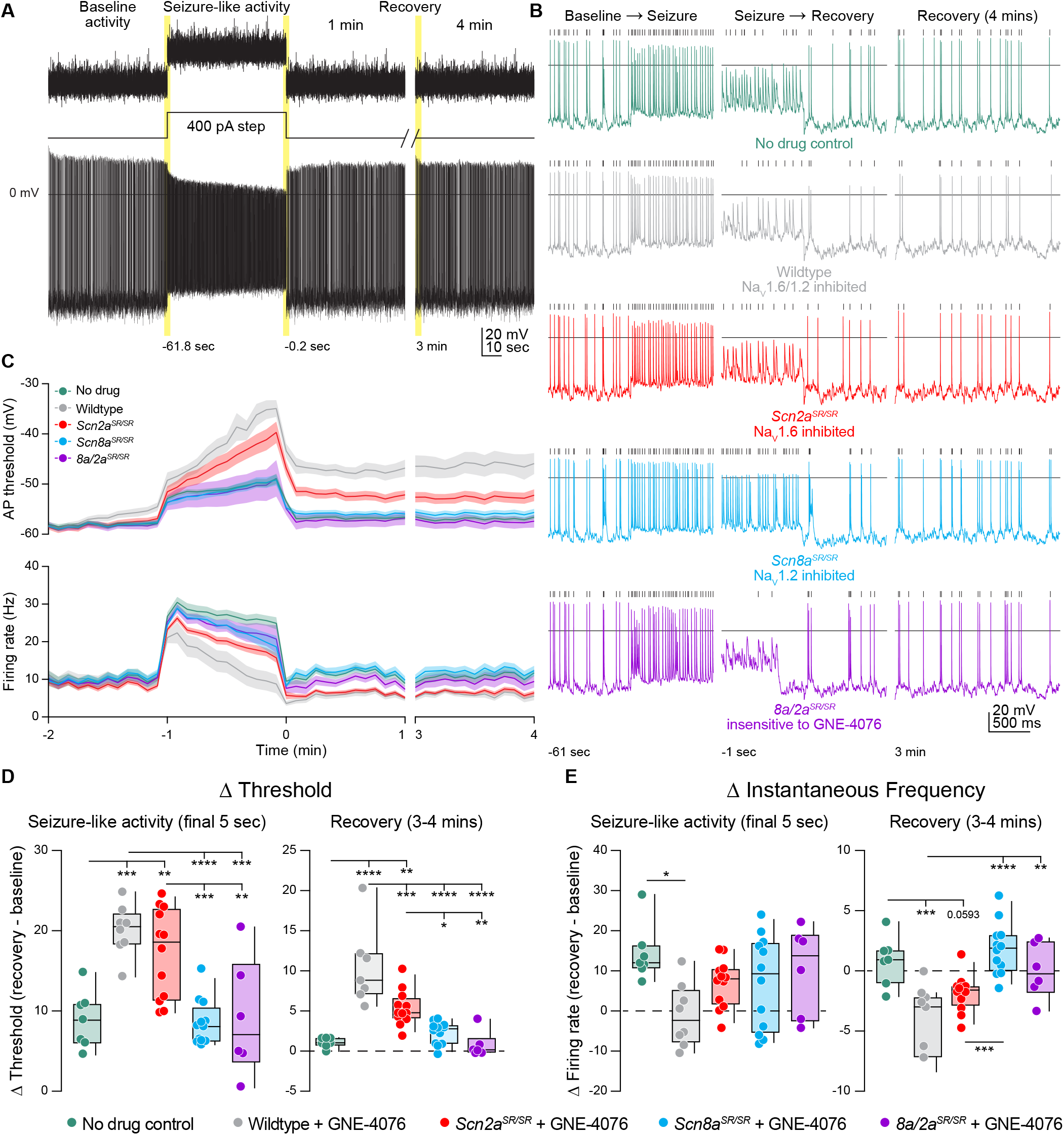
GNE-4076 onboarding following seizure-like activity continually impacts neuronal firing into recovery. (A) Stimulation protocol and example firing trace of cell injected with fluctuating post-synaptic potentials (PSPs) randomly generated using a Poisson probability distribution function for 60 sec. PSPs were continuously applied to acquire baseline activity, seizure-like activity and recovery activity. During seizure-like activity, a 400 pA step was applied in addition to the PSP. Recovery was continuously recorded for up to 4 mins post seizure-like activity. (B) Zoomed-in example traces for all genotypes at the Baseline → Seizure transition, Seizure → Recovery transition and start of 3-4 min recovery period (highlighted in (A)). Solid horizontal black bar represents membrane potential (V_m_) of 0 mV. Tick marks above traces represent detected spike defined as a change in V_m_ of 15 V/s or greater. (C) Threshold (mV) or instantaneous firing frequency (Hz) binned in 5 sec increments for all genotypes in (B). Solid lines represent mean value ± SEM. Timescale on x-axis mirrors activity presented in (A). (D) Summary of threshold data for the final 5 sec of seizure-like activity or entire 3-4 min recovery time-point in (C). Delta values are normalized to baseline activity (either at final 5 sec or entire period). Box plots show median and 90% tails. Circles represent individual cells (wildtype no drug, n=7; wildtype + GNE-4076, n=7-8; *Scn2a*^*SR/SR*^ + GNE-4076, n=12; *Scn8a*^*SR/SR*^ + GNE-4076, n=12; *8a/2a*^*SR/SR*^ + GNE-4076, n=6). One-way ANOVA, Holm-Šídák multiple comparisons test. *p<0.05, **p<0.01, ***p<0.001, ****p<0.0001. (E) Summary of instantaneous frequency data for the final 5 sec of seizure-like activity or entire 3-4 min recovery time-point in (C). Delta values are normalized to baseline activity (either at final 5 sec or entire period). Box plots show median and 90% tails. Circles represent individual cells (wildtype no drug, n=7; wildtype + GNE-4076, n=7-8; *Scn2a*^*SR/SR*^ + GNE-4076, n=12; *Scn8a*^*SR/SR*^ + GNE-4076, n=12; *8a/2a*^*SR/SR*^ + GNE-4076, n=6). One-way ANOVA, Holm-Šídák multiple comparisons test. *p<0.05, **p<0.01, ***p<0.001, ****p<0.0001.

In the presence of 200 nM GNE-4076, baseline firing rates were mostly stable across all conditions (Fig. S6A). Wildtype, no drug and *8a/2a*^*SR/SR*^ conditions observed no change in firing rate, while dual or selective Na_V_1.6 blockade saw a slight decrease by 60 sec of postsynaptic activity (Fig. S6B). Interestingly, we did see a slight increase in baseline firing by 60 sec when Na_V_1.2 was selectively targeted (Fig. S6A, B), furthering highlighting a paradoxical shift to hyperactive neurons when Na_V_1.2 availability decreases. By contrast, a slight depolarization in AP threshold was observed at baseline when both channels were sensitive to GNE-4076 (e.g., WT plus drug) and to a lesser extent with selective block of Na_V_1.6 (Fig. S6A, B). Given that this dose can completely block both Na_V_1.2 and Na_V_1.6 in HEK cells, a small amount of drug block was expected at baseline.

Seizure-like activity induction depolarized AP threshold markedly in all genotypes, and threshold depolarization persisted into the recovery phase for all neurons experiencing Na_V_1.6 or dual inhibition (Fig. 7C, D). Seizure-like activity increased spike rate across all genotypes at the onset of seizure-like activity induction (Fig. 7C, E). All cells exhibited some degree of AP accommodation during this stimulus, including drug-naïve controls and cases where both Na_V_1.2 and Na_V_1.6 were insensitive to GNE-4076 block. WT cells exposed to GNE-4076 accommodated markedly, returning to baseline firing rates at the end of the seizure-like stimulus (Fig. 7C, E). A less dramatic effect was noted when Na_V_1.6 was blocked alone (Fig. 7E). By contrast, cells where Na_V_1.2 could be blocked alone were not appreciably different than untreated controls (Fig. 7E). Furthermore, some cells that remained hyperactive with selective Na_V_1.2 block compared to controls cell throughout the recovery phase. Together, these data suggest that use-dependent pharmacology that targets Na_V_1.2 and Na_V_1.6 may be most beneficial if designed for higher potency at Na_V_1.6.

## DISCUSSION

Here, we combined genetic and pharmacological strategies to transiently, selectively and reversibly inhibit either Na_V_1.6 or Na_V_1.2 function using activity-dependent ASCs, which bind channels in the inactivated state. We show that acute blockade of either isoform has opposing effects on neuronal output: inhibition of Na_V_1.6 decreases AP output, whereas Na_V_1.2 increases AP output. Given this, we found that block of Na_V_1.6 rather than Na_V_1.2 was more effective at tempering spiking activity in cells driven to seizure-like levels of AP output. This suggests that pharmacology tuned to preferentially block Na_V_1.6 over Na_V_1.2 in a use-dependent manner may be useful as an anti-epileptic.

### Functional implications of differential compartmental Na_**V**_ isoform expression

Electrophysiological recordings of AP propagation delays between somatic and axonal recordings demonstrate that APs initiate ~35 to 50 microns from the soma, in the distal AIS^2,3,7,8^. This region is enriched with Na_V_ 1.6^8,9,44,58,59^. Consistent with this, we find that inhibition of Na_V_ 1.6 alone alters AP initiation, with an increase in AP threshold and decrease in total AP number from block of this isoform. Following distal AIS AP initiation, APs sequentially propagate across different neuronal compartments to influence local excitability^2,3,7^. Forward propagation was not examined here, but prior reports demonstrate that it is supported almost exclusively by Na_V_ 1.6 in nodes of Ranvier and boutons^9,13,14^. By contrast, backpropagation through the proximal AIS and soma recruits a mix of Na_V_1.6 and Na_V_1.2.

Prior work has suggested that Na_V_1.2 channels localized to the proximal AIS are critical for backpropagation, with modeling suggesting that APs would fail to effectively backpropagate into the soma in the absence of proximal AIS Na 1.2 channels^8^. While some efforts have been made to test this empirically with conditional *Scn2a* knockout, such experiments are imperfect, as Na_V_1.6 redistributes in the AIS of cells that lack Na_V_ 1.2^4^. Here, we were able to test the role of Na_V_ 1.2 in backpropagation without compensatory effects, using acute pharmacological inhibition. Backpropagation failures were not observed, consistent with results from conditional *Scn2a* knockout^4^. However, we did observe a reduction in the velocity at which the AIS component of the AP depolarized when Na_V_1.2 was blocked selectively (Fig. S3B-C). This suggests that proximal AIS-localized Na_V_1.2 do aid in boosting backpropagating APs as they transit from the distal AIS to the soma. Nevertheless, Na_V_1.2 channels are still dispensable for recruitment of somatic Na_V_ channels, at least in mouse mPFC layer 5b pyramidal cells. Whether similar effects are observed in other cells, including those with larger somata that may be more difficult to depolarize in the absence of Na_V_1.2, remains to be tested.

Na_V_1.2 expression in pyramidal cell dendrites appears to have two roles in neuronal excitability. Intuitively, these channels provide local inward current that boosts dendritic excitability, leading to a recruitment of voltage-gated calcium channels^5,6^. But this depolarization also appears to be critical for recruitment of dendritically localized voltage-gated potassium channels that contribute to AP repolarization and net membrane potential between spikes^4^. Indeed, conditional knockout of Na_V_ 1.2 paradoxically increases AP output, and we suggested previously that this was due in large part to loss of interactions between dendritic Na_V_ channels and K_V_ channels. But given observed changes in Na_V_1.6 function as well as potential for other compensatory changes in non Na_V_ ion channels, it was difficult to ascribe hyperexcitability purely to the loss of Na 1.2 alone ^50^. Here, we showed that identical increases in excitability—associated with depolarization of membrane potential between APs—could be observed with acute Na_V_1.2 block. This indicates that such effects can be due purely to the interplay between dendritic Na_V_1.2 and K_V_ channels, and that modifications to potassium channel distribution or function are not necessary for such effects.

Similar to Na_V_1.2 conditional knockout, where Na_V_1.6 is upregulated, knockout of Na_V_1.6 results in an increase in Na_V_1.2 expression, at least in the AIS. In pyramidal cells from mice constitutively lacking Na_V_1.6, Na_V_1.2 occupies the entirety of the AIS rather than just the region proximal to the soma^10,16,44^. Thus, it has not been possible to evaluate the role of Na_V_1.6 in its normal distribution using genetic manipulations. Here, we find that acute Na_V_1.6 block reduces AP output with a concomitant increase in AP threshold. This contrasted markedly with block of Na_V_1.2, which had no effect on threshold. Of note, this distinction can be leveraged to assess the specificity of other Na_V_-targeting pharmacology that has been suggested to have specificity at select isoforms, as any change in threshold indicates that drugs are interacting with Na_V_1.6^10,21^.

### ASC pharmacology for suppression of neuronal hyperexcitability

Epilepsy can arise from genetic and non-genetic factors, often with unexplained etiology^60,61^. Seizure onset is broadly classified as an electrical imbalance of cellular and network activity that favors hyperexcitability^62^, whether it be cell intrinsic, synaptic, or due to complex network effects^63,64^. Thus, seizure suppression can be targeted at multiple levels. Nevertheless, proper dosing can be difficult, as one aims to provide drug concentrations that temper excess activity but limit side effects like sedation associated with elevated drug concentrations.

For genetically defined sodium channelopathies, attention has been directed to neuronal cell-types that express each isoform (e.g., *SCN1A*: Na 1.1, *SCN2A*: Na 1.2 and *SCN8A*: Na 1.6)^65,66^. Seizures resulting from *SCN1A* loss of function limit inhibitory neuron excitability, thereby disinhibiting excitatory pyramidal cells^37,55,67^. *SCN2A* gain-of-function results in hyperexcitability in pyramidal cells, especially in early development when Na_V_1.2 channels are the sole isoform expressed in the AIS^12,68^. Later in development, *SCN8A*-encoded Na_V_ 1.6 channels are expressed more ubiquitously in the AIS of most cell classes, supporting AP initiation in both excitatory and inhibitory cells^37,53,59,69^.

With these cellular distributions and mechanisms of action, guidelines have emerged for treatment. For *SCN1A* loss-of-function, sodium channel blocking anti-epileptics are typically counter-indicated^70,71^. This is because currently prescribed sodium channel blockers are nonselective, with little preference for Na_V_ 1.1, 1.2 or 1.6^72^. Thus, further block of the remaining Na_V_1.1 leads to even more disinhibition of excitatory cells before any beneficial effects of blocking Na_V_ channels in excitatory cells are realized. Similarly, nonspecific sodium channel blockers are counter-indicated for *SCN2A* loss-of-function seizures, as they tend to increase seizure severity. Instead, non-selective Na_V_ inhibitors are useful when channel gain-of-function affects excitatory cells, as is the case for both *SCN2A* and *SCN8A* gain-of-function cases.

ASCs may be useful within each of these domains, as their chemistry can be adjusted to bias binding to specific isoforms in an activity-dependent manner^24–27^. Indeed, compounds similar to those used here that inhibit both Na_V_1.2 and Na_V_1.6 (but not other Na_V_ channels) are effective at suppressing chemoconvulsant-induced seizures in *ex vivo* models^25^. This parallels our observations, where excess activity was dampened most effectively by dual block of Na_V_1.2 and Na_V_1.6 (Fig. 6C). This effect appears due in large part to block of Na_V_1.6, since, ultimately, this is the isoform responsible for AP initiation. Consistent with this, Na_V_1.6-preferring ASCs show promise in protecting from chemoconvulsant-induced seizures *in vivo*^26,27^. Similar to our results, Johnson et al. also shows that a selective ASC for Na_V_1.6 (NBI-921352) preferentially targets cortical pyramidal cells, unlike Carbamazepine that non-selectively targets many Na_V_ isoforms leading to impaired interneuron activity^27^. In fact, drug therapies using agents like Carbamazepine, Lamotrigine and Phenytoin that have little to no selectivity for Na_V_ isoforms can have adverse effects in certain situations^62^. Thus, it may be that preferential block of Na_V_1.6 would be beneficial in multiple sodium channelopathy conditions, including those associated with channel loss, since network hyperexcitability is ultimately dictated by Na_V_1.6 function in the AIS of excitatory neurons after the first months of life.

Beyond selectivity, there may also be advantages to ASC activity-dependent properties. The ASC binding pocket is hidden from drug in a channel’s closed state. This feature, combined with on- and off-rate kinetics, results in an accumulation of block primarily for highly-active neurons (Figs. 4, 6). Similar to on-demand, closed-loop electrical or optogenetic approaches for seizure intervention^73–75^, ASCs are essentially biased to suppress prolonged, high-frequency activity, including the type of activity commonly observed during seizures. In this study, our focus centered mainly on seizure-like activity across individual neurons. Future studies can extend this approach to examining networks *ex vivo* and systems *in vivo*.

## ACKNOWLEDGEMENTS

We thank Drs. JP Johnson, Natali Minassian, Fiona Scott, and members of the Bender Lab for extensive discussions related this work. This work was supported by NIH grants K00 MH134674 (JDG), MH125978 and MH126960 (KJB), by the Hartwell foundation through an Individual Biomedical Research Award (RBS) and by FamiliesSCN2A through an Action Potential award (RBS).

## METHODS

### RESOURCES AVAILABILITY

#### Lead contact

Any additional information or enquires related to reagents or resources should be directed to the lead contact, Kevin J. Bender (kevin.bender@ ucsf.edu).

#### Materials availability

The transfer of unique reagents generated for this study will be made available upon request. A Materials Transfer Agreement may be required.

#### Data and code availability

This study did not generate any unique datasets or code. Data reported here will be made available by lead contact upon reasonable request.

### EXPERIMENTAL MODELS AND SUBJECTS DETAILS

#### Mouse strains

All animal procedures are in accordance with the Institutional Animal Care and Use Committee (IACUC) guidelines in accordance with the University of California, San Francisco (UCSF). The following mouse strains were used in this study: C57BL/6J, YW−≥SR Na_V_ 1.2 KI (*Scn2a*^*SR/ SR*^), YW->SR Na_V_ 1.6 KI (*Scn8a*^*SR/SR*^) and YW−≥SR dual KI (*8a/2a*^*SR/SR*^). All experimental procedures were performed on mice maintained in-house on a 12:12 hour light-dark cycle under standard conditions with *ad libitum* access to food and water. For genotyping, genomic DNA was isolated from tail clip biopsies for PCR. Both male and female mice aged postnatal day (P)18-59 were used across all genotypes. C57BL/6J mice were obtained from Jackson Laboratories and YW−≥SR KI mice were developed by the Hackos Lab (Genentech).

#### Generation of *SCN8A* or *SCN2A* YW->SR KI mouse

As described previously^29^, CRISPR/Cas9 technology^30,31^ was used to generate a genetically modified mouse strain with either an *Scn8a* or *Scn2a* YW−≥SR knock-in mutation. A single guide RNA (sgRNA) target and protospacer adjacent motifs (PAM) were identified for *Scn8a* ENSMUSG00000023033 or *Scn2a* ENSMUSG00000075318 genomic regions of interest using the CRISPR design tool (Benchling) that uses the algorithm described by Hsu et al.^32^ to provide ‘MIT’ specificity scores for each sgRNA, as well as the top 15 predicted off-target loci and corresponding MIT off-target scores. The same guide target and PAM were used for both genes. Guide target: 5’ CATTCTCTACTGGATTAATC 3’; PAM: TGG with an algorithm score of 42.3.

For the YW−≥SR mutation on the *Scn8a* gene, predicted cut sites are between 100,933,463-100,933,464 genome coordinates. The following oligonucleotide donor sequence was used: 5’ ATGCTTATCTGCCTTAACATGGTGACCATGATGGTGGAGACAGACACACAGAGCAAGCAGATGGAGAACATTCTCTCTCGGATTAATCTGGTCTTCGTCATCTTCTTCACCTGCGAGTGTGTGCTCAAAATGTTTGCCTTGAGACACTACTATTTC 3’. The first point mutation of Y1553S (TAC->TCT) is located at 100,933,454-100,933,456 genome coordinates, and a second point mutation of W1554R (TGG->CGG) is located at 100,933,457-100,933,459 genome coordinates.

For the YW−≥SR mutation on the *Scn2a* gene, predicted cut sites are between 166,900-166,901 genome coordinates. The following oligonucleotide donor sequence was used: 5’ GAAATAGTAGTGTCTCAAGGCAAACATTTTGAG CACACACTCGCAGGTGAAGAAGATGACGAAGACCAGGTTGATCCGA GAGAGAATGTTCTCCATCTGCTTGCTCTGTGTGTCTGTCTCCACCATC ATGGTCACCATGTTAAGGCAGATAAGCAT 3’. The first point mutation of Y1553S (TAC->TCT) is located at 101035573-101035575 genome coordinates, and a second point mutation of W1554R (TGG->CGG) is located at 101035576-101035578 genome coordinates. Additionally, two silent mutations were created in the beginning of the gRNA to prevent Cas9 from cutting the donor oligo. The first silent mutation (ATT->ATC) is located at 101035579-101035581 genome coordinates and the second silent mutation (AAT->AAC) 101035582-101035584 genome coordinates.

After homology-directed repair of Cas9-induced chromosome breaks with the oligonucleotide donor, the YW−≥SR protein will be expressed. Once a sgRNA decision was finalized, the off-target list was used to identify the top 15, and next-generation sequencing (NGS) amplicon primers were designed for the on-target locus, and each of the off-targets synthetic guide RNA was obtained from Synthego. CAS9 protein was obtained from PROTEIN SOURCE and complexed with sgRNA before microinjection. Reagent concentrations for microinjection were as follows: 25 ng/μl Cas9 mRNA (Thermo Fisher; A29378) + 13 ng/μl sgRNA (Synthego), Oligonucleotide donor (50 ng/μl) (IDT).

After zygote microinjection and embryo transfer, genomic DNA was prepared from tail tip biopsies of potential G0 founders, and G0 animals were first analyzed by droplet digital PCR^33^ (Bio-Rad). Primers were used to amplify the HDR event (ON Target) and the 15 most likely off-target sites. Only G0 mosaic founders positive for the intended mutation were screened by targeted amplicon NGS. Amplicons were submitted for NGS analysis.

Founders were selected for mating with wild-type C57BL/6N mice for germline transmission of the gene edited chromosome. Subsequent analysis of genomic DNA from G1 pups was used to confirm germline transmission of the targeted gene and the absence of off-target hits elsewhere in the genome.

To generate the dual *8a/2a*^*SR/SR*^ line, *Scn2a*^*SR/SR*^ and *Scn8a*^*SR/SR*^ mice were crossed and analysis of genomic DNA was used to confirm germline transmission of the targeted gene.

### METHOD DETAILS

#### Synthesis of GNE4076

GNE-4076 was synthesized as previously described in Roecker et al.^28^, and is is Compound 5 in original manuscript. We use GNE-4076 throughout this manuscript.

#### Generation of *SCN2A* YW->SR and SCN8A YW->SR constructs

The double mutation Y1564S/W1565R was introduced into the adult splice isoform of recombinant human Na_V_1.2 (NCBI accession number NM_021007; AddGene #162279) using site directed mutagenesis as previously described^34^. The corresponding mutations (Y1555S/W15556R) were engineered in recombinant human Na_V_1.6 (adult isoform) as previously described^35^. Mutagenic primer sequences are presented in Table 1. All plasmids were nanopore sequenced (Primoridium Labs, Arcadia, CA) to confirm the variants and exclude unwanted mutations.

#### Electrophysiology in immortalized cell lines

HEK293 cells were transfected with WT or mutant Na_V_1.2 using the Invitrogen Lipofectamine LTX kit, whereas Na_V_1.6 plasmids were electroporated into the LoNav derivative of ND7/23 cells using Maxcyte technology as previously described^35^.

Na_V_ channel currents were recorded from HEK293 or ND7/23 cells using whole-cell patch clamp using a Molecular Devices Axopatch 200B amplifier. The recording pipet intracellular solution contained (in mM): 120 CsF, 10 NaCl, 2 MgCl_2_, 10 HEPES, adjusted to pH 7.2 with CsOH. The extracellular recording solution contained (in mM): 155 NaCl, 3 KCl, 1 MgCl_2_, 1.5 CaCl_2_, 10 HEPES, adjusted to pH 7.4 with NaOH. Currents were recorded at 20 kHz sampling frequency and filtered at 5 kHz. Series resistance compensation was applied at 80%. Solutions containing GNE-4076 were applied using a Fluicell Dynaflow perfusion system.

We characterized dose response curves in both WT and mutant Na_V_1.2 and Na_V_1.6 channels by pulsing cells from −80 mV to 0 mV for 20 msec at a rate of 0.5 Hz to establish a baseline current. Cells were then perfused with GNE-4076 starting at 30 nM and increased up to 100 μM. To allow adequate GNE-4076 onboarding to NaV isoforms/mutants at each successive dose, cells were depolarized to 0 mV for 10 sec followed by similar test pulses used to acquire baseline current. Between dose increases, cells were held at −80 mV to allow adequate unbinding of GNE-4076.

We also characterized the activation and steady-state inactivation properties of both WT and mutant Na_V_1.2 and Na_V_1.6 channels. To measure activation, we used a holding voltage of −80 mV, a short 20 ms pre-pulse to −120 mV, and a 30 ms pulse to voltages ranging from −100 mV to −20 mV in steps of 5mV at a rate of once per 3 sec. P/4 leak subtraction was used to reduce leak currents. The peaks of the resulting Na_V_ currents were measured. Conductance was calculated using the equation G = I / (V – V_Na_), normalized, and plotted as a function of voltage. To measure inactivation, we started at a holding voltage of −120 mV to bring the channels fully into the closed state (about 1 min). We then pulsed to 0 mV while reducing the holding voltage from −120 mV to +30 mV in steps of 5 mV at a rate of once per 5 sec. The peaks of the resulting Na_V_ currents were measured, normalized and plotted as a function of voltage. To quantify these biophysical properties, we fit the activation and inactivation curves to the following Boltzmann equations:

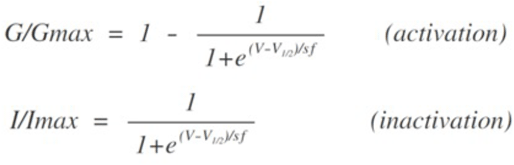

V_1/2_ is the voltage where half-maximal peak conductance (or current) was observed, and sf is the slope factor.

#### *Ex vivo* electrophysiology

Mice aged P18-P59 were anesthetized with isoflurane prior to harvesting the brain. Dissected brains were immediately placed in cutting solution (4°C) containing 87 mM NaCl, 25 mM NaHCO_3_, 25 mM glucose, 75 mM sucrose, 2.5 mM KCl, 1.25 mM NaH_2_PO_4_, 0.5 mM CaCl_2_ and 7 mM MgCl_2_ that is bubbled with 5% CO_2_ / 95% O_2_. Coronal slices were prepared from the medial prefrontal cortex (PFC) at a thickness of 250 microns and placed in a holding chamber warmed to 33C for 30 minutes. Slices were then allowed to recover at room temperature until recording. Recordings were performed at 31-33°C in a solution containing 125 mM NaCl, 2.5 mM KCl, 2 mM CaCl_2_, 1 mM MgCl_2_, 25 mM NaHCO_3_, 1.25 mM NaH_2_PO_4_ and 25 mM glucose that is bubbled with 5% CO_2_ / 95% O_2_. Osmolarity of the recording solution was adjusted to approximately 309 mOsm.

Neurons were identified using differential interference contrast (DIC) optics for conventional visually guided whole-cell recordings. For current clamp experiments, pipettes were pulled from Schott 8250 glass, with a tip resistance of 3-4 M, and filled with a K-gluconate based internal solution containing 113 mM K-Gluconate, 9 mM HEPES, 4.5 mM MgCl_2_, 0.1 mM EGTA, 14 mM Tris_2_-phosphocreatine, 4 mM Na_2_-ATP, 0.3 mM Tris-GTP; 290 mOsm; pH: 7.2-7.25. All data were corrected for measured junction potentials of 12 mV.

All electrophysiology data were acquired through custom protocols generated in IgorPro (Wavemetrics) via Multiclamp 700A or 700B amplifiers (Molecular Devices). Action potential (AP) waveform measurements were acquired at 50 kHz and low-pass Bessel filtered at 20 kHz for all experiments except for data presented in Fig. 6 (acquired at 10 kHz and filtered at 3 kHz). Pipette capacitance was compensated in all current-clamp recordings to 50% of the fast capacitance measured after membrane seals were established in voltage-clamp. The bridge was balanced, and series resistance was kept < 18 MΩ in all recording. Any cells with input resistance changes exceeding ± 15% were omitted from dataset. A quartz electrode holder (Sutter Instrument) was used to collect all recording to minimize physical drift of recording electrodes.

All recordings were made from medial prefrontal cortex (PFC) layer 5b, thick-tufted pyramidal tract (PT) neurons. Pyramidal neuron identity was confirmed by assessing membrane responses to hyperpolarizing current (−400 pA, 120 ms), with PT neurons defined as those that exhibit membrane depolarization overshoot that peaks withing 90 ms of current step offset^36^. AP threshold, AIS dV/dt and peak dV/dt measurements were determined from either the first AP or all APs within a spike train evoked by a stimulus. AP threshold was defined as the membrane potential (V_m_) when the AP speed (dV/dt) exceeds 15 V/s. Somatic peak (V/s) was defined as the max value in the first derivative (dV/dt) of an AP. AIS max (V/s) was defined as the trough or saddle point in the second derivative (d^2^V/dt^2^) that occurs during the depolarizing phase of an AP before the somatic peak^37^. AIS inflection point was defined as the peak of the second derivative (d^2^V/ dt^2^) that occurs prior to the AIS max or somatic peak. Frequency was defined as the spike number in a train elicited per second (spikes/sec or Hz). Afterhyperpolarization (AHP) was defined as the minimum voltage between 2 spikes in a train.

Spike trains were evoked by injecting current (250-350 pA, 10 sec or 300 ms duration) in the presence or absence of 200 nM GNE-4076. In recovery experiments, neurons were held at −12 mV for 30 sec (voltage-clamp) to depolarize cells, activate Na_V_ channels and expose the ASC binding pocket to maximize GNE-4076 onboarding and Na_V_ inhibition. The interstimulus interval given during the recovery period was as follows: 2 sec, ~20-30 sec duration; 5 sec, ~30-45 sec duration; 15 sec, ~60-90 sec duration; 30 sec, ~120-180 sec duration; and 60 sec, ~60-240 sec duration. In experiments mimicking cellular activity, a post-synaptic potential train was randomly generated using a Poisson probability distribution function for 60 sec with a frequency of 50 Hz and amplitude standard deviation of 200 pA.

For nucleated patch voltage-clamp experiments, recordings were made at 31-33°C in a solution containing 125 mM NaCl, 2.5 mM KCl, 2 mM CaCl_2_, 1 mM MgCl_2_, 25 mM NaHCO_3_, 1.25 mM NaH_2_PO_4_, 15 mM glucose, 4 mM TEA, 1 mM 4-AP, and 10 μM nifedipine, with or without 200 nM GNE-4076, (bubbled with 5% CO_2_ / 95% O_2_). A Cs-methanesulfonate based recording solution was used: 110 mM CsMeSO_3_, 40 mM HEPES, 1 mM KCl, 4 mM NaCl, 4 mM Mg-ATP, 10 mM Na-phosphocreatine, 0.4 mM Na_2_-GTP, and 0.1 mM EGTA; 290 mOsm; pH: 7.2-7.25. Whole-cell recordings were established before withdrawing pipettes from the slice, pulling a region of the somatic membrane with the pipette. Neurons were held to −80 mV and stepped to −12 mV every 2 sec, 5-10 times, then neurons were held to −12 mV for 10 sec to onboard GNE-4076, then returned to −80 mV and probed for Na_V_ recovery with steps to −12 mV every 2 sec. Leak currents were subtracted with a P/8 protocol using steps from −80 to −90 mV.

#### Compartmental modeling

A compartmental model was constructed within the NEURON environment to simulate a layer 5 pyramidal neuron as described before^4,38,39^. A multi-compartmental model, originally developed by the Blue Brain Project, was implemented to reflect the morphology detailed by Ramaswamy and Markram^40^ and electrophysiological features were adjusted to reflect empirically obtained data^4,38^. We modified the model by replacing the aggregated sodium conductances (NaT and NaP) with distinct Na_V_1.2 and Na_V_1.6 channels to match empirically observed distributions^8^. Na 1.2 and Na 1.6 channels were distributed throughout the cell with equal levels in the soma and 20 μm of the proximal dendrites. Na_V_1.2 was solely expressed in dendrites more distal to the soma, inferred based on AP-evoked sodium imaging observations in *Scn2a*^*+/-*^ conditions^6^.

The two axonal compartments were subdivided into an axon initial segment and distal axon. Within the AIS, Na_V_1.2 and Na_V_1.6 were distributed with increased Na_V_1.2 in the proximal AIS and increased Na_V_1.6 in the distal AIS to recapitulate the channel distribution as previously observed empirically^8^. Na_V_ 1.2 was not included in the distal AIS or axon where only Na_V_1.6 is present, including an enriched region to model a node of Ranvier. To simulate blocking of Na_V_1.2 and Na_V_1.6, each channel’s density was globally reduced in 10% increments from 100% to 0%. Both Na_V_1.2 and Na_V_ 1.6 channels were represented using the Hodgkin-Huxley formalism^41^. Parameter optimization for both channels was conducted using an evolutionary algorithm from BluePyOpt^42^ and adapted for use with the computational resources at the National Energy Research Computing Center similar to Ladd et al.^43^.

To validate parameter sensitivity, we constructed multiple variations of our model with different distributions of Na_V_1.2 and Na_V_1.6 in the AIS. The crossover point at which the Na_V_1.2 and Na_V_1.6 distribution curves intersect was shifted distally in increments of 3.75 microns to make 5 variations of the AIS with increasing density of Na_V_1.2. With the wildtype crossover point located 15 microns distal to the soma, the 5 right shifted variations have crossover points at 18.75 μm, 22.5 μm, 26.25 μm, 30 μm, and 33.75 μm respectively. Two variations of the AIS with increased Na_V_1.6 density were created by shifting the crossover point proximally in 3.75 micron increments to 11.25 μm and 7.5 μm. To see how our model would behave with different AIS sodium channel distributions, we varied the ratio of Na_V_1.2: Na_V_1.6 density in the whole cell to test different conditions at each AIS crossover point and measured the resulting threshold, AIS peak, and somatic peak of the phase plane plot. Na_V_1.2 percentage was decreased from 90% to 0% in 10% increments as Na_V_1.6 percentage was concurrently increased from 10% to 100% in 10% increments. The Na_V_1.2: Na_V_1.6 ratio of 100%:0% did not produce any action potentials and was therefore not included in the analysis. In instances where the somatic portion of the phase plane was not a true peak that consisted of adjacent values less than a local maxima, the prominence was estimated by taking the index of the second derivative value closest to zero and calculating the dV/dt value at that index.

### QUANTIFICATION AND STATISTICAL ANAYLYSIS

Data are reported as absolute values or the absolute difference from baseline (delta, Δ). For Δ values, data were normalized either to the initial 500 ms of the stimulus (Fig. 2–3) or baseline spiking (Fig. 4–5). Time course graphs are represented as a mean ± standard error. Summary graphs are represented with boxplots showing the median, quartiles and 90% tails or with violin plots. All summary graphs overlay individual datapoints and represent recordings from single cells (reported n) for all electrophysiology experiments. Data were obtained from 5-9 animals (both sexes) per condition, which are standard group sample sizes used in the field. Analysis was performed blind to genotype ± drug. Statistical analysis was performed using Prism 10 (Graphpad Software). Quantified mean ± standard error and statistical test used is noted in figure legends. Significance was set at an alpha value of 0.05 and ‘ns’ indicates no significance.

#### Author Contributions

DHH and KJB conceived the project. JDG, CW, RA, EB, TF, JMD, TVA, ALG, RBS, DDH and KJB designed and performed experiments. JDG, CW, RA, EB, TF, JMD, TVA, ALG, RBS, DDH and KJB analyzed the data. JDG and KJB wrote the manuscript. EB, TF, ALG, RBS, and DDH reviewed and edited the manuscript.

## REFERENCES

1. Meeks, J.P., and Mennerick, S. (2007). Action potential initiation and propagation in CA3 pyramidal axons. J Neurophysiol 97, 3460–3472. 10.1152/jn.01288.2006.

2. Shu, Y., Duque, A., Yu, Y., Haider, B., and McCormick, D.A. (2007). Properties of action-potential initiation in neocortical pyramidal cells: evidence from whole cell axon recordings. J Neurophysiol 97, 746–760. 10.1152/jn.00922.2006.

3. Palmer, L.M., and Stuart, G.J. (2006). Site of action potential initiation in layer 5 pyramidal neurons. J Neurosci 26, 1854–1863. 10.1523/jneurosci.4812-05.2006.

4. Spratt, P.W.E., Alexander, R.P.D., Ben-Shalom, R., Sahagun, A., Kyoung, H., Keeshen, C.M., Sanders, S.J., and Bender, K.J. (2021). Paradoxical hyperexcitability from Na(V)1.2 sodium channel loss in neocortical pyramidal cells. Cell Rep 36, 109483. 10.1016/j.celrep.2021.109483.

5. Spratt, P.W.E., Ben-Shalom, R., Keeshen, C.M., Burke, K.J., Jr., Clarkson, R.L., Sanders, S.J., and Bender, K.J. (2019). The Autism-Associated Gene Scn2a Contributes to Dendritic Excitability and Synaptic Function in the Prefrontal Cortex. Neuron 103, 673-685.e675. 10.1016/j.neuron.2019.05.037.

6. Nelson, A.D., Catalfio, A.M., Gupta, J.P., Min, L., Caballero-Florán, R.N., Dean, K.P., Elvira, C.C., Derderian, K.D., Kyoung, H., Sahagun, A., et al. (2024). Physical and functional convergence of the autism risk genes Scn2a and Ank2 in neocortical pyramidal cell dendrites. Neuron 112, 1133-1149.e1136. 10.1016/j.neuron.2024.01.003.

7. Baranauskas, G., David, Y., and Fleidervish, I.A. (2013). Spatial mismatch between the Na+ flux and spike initiation in axon initial segment. Proc Natl Acad Sci U S A 110, 4051–4056. 10.1073/pnas.1215125110.

8. Hu, W., Tian, C., Li, T., Yang, M., Hou, H., and Shu, Y. (2009). Distinct contributions of Na(v)1.6 and Na(v)1.2 in action potential initiation and backpropagation. Nat Neurosci 12, 996–1002. 10.1038/nn.2359.

9. Tian, C., Wang, K., Ke, W., Guo, H., and Shu, Y. (2014). Molecular identity of axonal sodium channels in human cortical pyramidal cells. Front Cell Neurosci 8, 297. 10.3389/fncel.2014.00297.

10. Ye, M., Yang, J., Tian, C., Zhu, Q., Yin, L., Jiang, S., Yang, M., and Shu, Y. (2018). Differential roles of Na(V)1.2 and Na(V)1.6 in regulating neuronal excitability at febrile temperature and distinct contributions to febrile seizures. Sci Rep 8, 753. 10.1038/s41598-017-17344-8.

11. Yu, Y., Shu, Y., and McCormick, D.A. (2008). Cortical action potential backpropagation explains spike threshold variability and rapid-onset kinetics. J Neurosci 28, 7260–7272. 10.1523/jneurosci.1613-08.2008.

12. Jenkins, P.M., and Bender, K.J. (2024). Axon Initial Segment Structure and Function in Health and Disease. Physiol Rev. 10.1152/physrev.00030.2024.

13. Caldwell, J.H., Schaller, K.L., Lasher, R.S., Peles, E., and Levinson, S.R. (2000). Sodium channel Na(v)1.6 is localized at nodes of ranvier, dendrites, and synapses. Proc Natl Acad Sci U S A 97, 5616–5620. 10.1073/pnas.090034797.

14. Kaplan, M.R., Cho, M.H., Ullian, E.M., Isom, L.L., Levinson, S.R., and Barres, B.A. (2001). Differential control of clustering of the sodium channels Na(v)1.2 and Na(v)1.6 at developing CNS nodes of Ranvier. Neuron 30, 105–119. 10.1016/s0896-6273(01)00266-5.

15. Fleidervish, I.A., Lasser-Ross, N., Gutnick, M.J., and Ross, W.N. (2010). Na+ imaging reveals little difference in action potential-evoked Na+ influx between axon and soma. Nat Neurosci 13, 852–860. 10.1038/nn.2574.

16. Katz, E., Stoler, O., Scheller, A., Khrapunsky, Y., Goebbels, S., Kirchhoff, F., Gutnick, M.J., Wolf, F., and Fleidervish, I.A. (2018). Role of sodium channel subtype in action potential generation by neocortical pyramidal neurons. Proc Natl Acad Sci U S A 115, E7184–e7192. 10.1073/pnas.1720493115.

17. Zhang, J., Chen, X., Eaton, M., Wu, J., Ma, Z., Lai, S., Park, A., Ahmad, T.S., Que, Z., Lee, J.H., et al. (2021). Severe deficiency of the voltage-gated sodium channel Na(V)1.2 elevates neuronal excitability in adult mice. Cell Rep 36, 109495. 10.1016/j.celrep.2021.109495.

18. Ragsdale, D.S., and Avoli, M. (1998). Sodium channels as molecular targets for antiepileptic drugs. Brain Res Brain Res Rev 26, 16–28. 10.1016/s0165-0173(97)00054-4.

19. Denomme, N., Lukowski, A.L., Hull, J.M., Jameson, M.B., Bouza, A.A., Narayan, A.R.H., and Isom, L.L. (2020). The voltage-gated sodium channel inhibitor, 4,9-anhydrotetrodotoxin, blocks human Na(v)1.1 in addition to Na(v)1.6. Neurosci Lett 724, 134853. 10.1016/j.neulet.2020.134853.

20. Bosmans, F., Rash, L., Zhu, S., Diochot, S., Lazdunski, M., Escoubas, P., and Tytgat, J. (2006). Four novel tarantula toxins as selective modulators of voltage-gated sodium channel subtypes. Mol Pharmacol 69, 419–429. 10.1124/mol.105.015941.

21. Filipis, L., Blömer, L.A., Montnach, J., Loussouarn, G., De Waard, M., and Canepari, M. (2023). Nav1.2 and BK channel interaction shapes the action potential in the axon initial segment. J Physiol 601, 1957–1979. 10.1113/jp283801.

22. Catterall, W.A., Goldin, A.L., and Waxman, S.G. (2005). International Union of Pharmacology. XLVII. Nomenclature and structure-function relationships of voltage-gated sodium channels. Pharmacol Rev 57, 397–409. 10.1124/pr.57.4.4.

23. de Lera Ruiz, M., and Kraus, R.L. (2015). Voltage-Gated Sodium Channels: Structure, Function, Pharmacology, and Clinical Indications. J Med Chem 58, 7093–7118. 10.1021/jm501981g.

24. Ahuja, S., Mukund, S., Deng, L., Khakh, K., Chang, E., Ho, H., Shriver, S., Young, C., Lin, S., Johnson, J.P., Jr., et al. (2015). Structural basis of Nav1.7 inhibition by an isoform-selective small-molecule antagonist. Science 350, aac5464. 10.1126/science.aac5464.

25. Goodchild, S.J., Shuart, N.G., Williams, A.D., Ye, W., Parrish, R.R., Soriano, M., Thouta, S., Mezeyova, J., Waldbrook, M., Dean, R., et al. (2024). Molecular Pharmacology of Selective Na(V)1.6 and Dual Na(V)1.6/Na(V)1.2 Channel Inhibitors that Suppress Excitatory Neuronal Activity Ex Vivo. ACS Chem Neurosci 15, 1169–1184. 10.1021/acschemneuro.3c00757.

26. Johnson, J.P., Jr., Focken, T., Karimi Tari, P., Dube, C., Goodchild, S.J., Andrez, J.C., Bankar, G., Burford, K., Chang, E., Chowdhury, S., et al. (2024). The contribution of Na(V)1.6 to the efficacy of voltage-gated sodium channel inhibitors in wild type and Na(V)1.6 gain-of-function (GOF) mouse seizure control. Br J Pharmacol 181, 3993–4011. 10.1111/bph.16481.

27. Johnson, J.P., Focken, T., Khakh, K., Tari, P.K., Dube, C., Goodchild, S.J., Andrez, J.C., Bankar, G., Bogucki, D., Burford, K., et al. (2022). NBI-921352, a first-in-class, Na(V)1.6 selective, sodium channel inhibitor that prevents seizures in Scn8a gain-of-function mice, and wild-type mice and rats. Elife 11. 10.7554/eLife.72468.

28. Roecker, A.J., Egbertson, M., Jones, K.L.G., Gomez, R., Kraus, R.L., Li, Y., Koser, A.J., Urban, M.O., Klein, R., Clements, M., et al. (2017). Discovery of selective, orally bioavailable, N-linked arylsulfonamide Na(v)1.7 inhibitors with pain efficacy in mice. Bioorg Med Chem Lett 27, 2087–2093. 10.1016/j.bmcl.2017.03.085.

29. Deng, L., Dourado, M., Reese, R.M., Huang, K., Shields, S.D., Stark, K.L., Maksymetz, J., Lin, H., Kaminker, J.S., Jung, M., et al. (2023). Nav1.7 is essential for nociceptor action potentials in the mouse in a manner independent of endogenous opioids. Neuron 111, 2642-2659.e2613. 10.1016/j.neuron.2023.05.024.

30. Cong, L., Ran, F.A., Cox, D., Lin, S., Barretto, R., Habib, N., Hsu, P.D., Wu, X., Jiang, W., Marraffini, L.A., and Zhang, F. (2013). Multiplex genome engineering using CRISPR/Cas systems. Science 339, 819–823. 10.1126/science.1231143.

31. Mali, P., Yang, L., Esvelt, K.M., Aach, J., Guell, M., DiCarlo, J.E., Norville, J.E., and Church, G.M. (2013). RNA-guided human genome engineering via Cas9. Science 339, 823–826. 10.1126/science.1232033.

32. Hsu, P.D., Scott, D.A., Weinstein, J.A., Ran, F.A., Konermann, S., Agarwala, V., Li, Y., Fine, E.J., Wu, X., Shalem, O., et al. (2013). DNA targeting specificity of RNA-guided Cas9 nucleases. Nat Biotechnol 31, 827–832. 10.1038/nbt.2647.

33. Hindson, B.J., Ness, K.D., Masquelier, D.A., Belgrader, P., Heredia, N.J., Makarewicz, A.J., Bright, I.J., Lucero, M.Y., Hiddessen, A.L., Legler, T.C., et al. (2011). High-throughput droplet digital PCR system for absolute quantitation of DNA copy number. Anal Chem 83, 8604–8610. 10.1021/ac202028g.

34. Thompson, C.H., Potet, F., Abramova, T.V., DeKeyser, J.M., Ghabra, N.F., Vanoye, C.G., Millichap, J.J., and George, A.L. (2023). Epilepsy-associated SCN2A (NaV1.2) variants exhibit diverse and complex functional properties. J Gen Physiol 155. 10.1085/jgp.202313375.

35. Vanoye, C.G., Abramova, T.V., DeKeyser, J.M., Ghabra, N.F., Oudin, M.J., Burge, C.B., Helbig, I., Thompson, C.H., and George, A.L., Jr. (2024). Molecular and cellular context influences SCN8A variant function. JCI Insight 9. 10.1172/jci.insight.177530.

36. Clarkson, R.L., Liptak, A.T., Gee, S.M., Sohal, V.S., and Bender, K.J. (2017). D3 Receptors Regulate Excitability in a Unique Class of Prefrontal Pyramidal Cells. J Neurosci 37, 5846–5860. 10.1523/jneurosci.0310-17.2017.

37. Favero, M., Sotuyo, N.P., Lopez, E., Kearney, J.A., and Goldberg, E.M. (2018). A Transient Developmental Window of Fast-Spiking Interneuron Dysfunction in a Mouse Model of Dravet Syndrome. J Neurosci 38, 7912–7927. 10.1523/jneurosci.0193-18.2018.

38. Hallermann, S., de Kock, C.P., Stuart, G.J., and Kole, M.H. (2012). State and location dependence of action potential metabolic cost in cortical pyramidal neurons. Nat Neurosci 15, 1007–1014. 10.1038/nn.3132.

39. Quinn, S., Zhang, N., Fenton, T.A., Brusel, M., Muruganandam, P., Peleg, Y., Giladi, M., Haitin, Y., Lerche, H., Bassan, H., et al. (2024). Complex biophysical changes and reduced neuronal firing in an SCN8A variant associated with developmental delay and epilepsy. Biochim Biophys Acta Mol Basis Dis 1870, 167127. 10.1016/j.bbadis.2024.167127.

40. Ramaswamy, S., and Markram, H. (2015). Anatomy and physiology of the thick-tufted layer 5 pyramidal neuron. Front Cell Neurosci 9, 233. 10.3389/fncel.2015.00233.

41. Hodgkin, A.L., and Huxley, A.F. (1952). A quantitative description of membrane current and its application to conduction and excitation in nerve. J Physiol 117, 500–544. 10.1113/jphysiol.1952.sp004764.

42. Van Geit, W., Gevaert, M., Chindemi, G., Rössert, C., Courcol, J.D., Muller, E.B., Schürmann, F., Segev, I., and Markram, H. (2016). BluePyOpt: Leveraging Open Source Software and Cloud Infrastructure to Optimise Model Parameters in Neuroscience. Front Neuroinform 10, 17. 10.3389/fninf.2016.00017.

43. Ladd, A., Kim, K.G., Balewski, J., Bouchard, K., and Ben-Shalom, R. (2022). Scaling and Benchmarking an Evolutionary Algorithm for Constructing Biophysical Neuronal Models. Front Neuroinform 16, 882552. 10.3389/fninf.2022.882552.

44. Tukker, A.M., Vrolijk, M.F., van Kleef, R., Sijm, D., and Westerink, R.H.S. (2023). Mixture effects of tetrodotoxin (TTX) and drugs targeting voltage-gated sodium channels on spontaneous neuronal activity in vitro. Toxicol Lett 373, 53–61. 10.1016/j.toxlet.2022.11.005.

45. Thouta, S., Waldbrook, M.G., Lin, S., Mahadevan, A., Mezeyova, J., Soriano, M., Versi, P., Goodchild, S.J., and Parrish, R.R. (2022). Pharmacological determination of the fractional block of Nav channels required to impair neuronal excitability and ex vivo seizures. Front Cell Neurosci 16, 964691. 10.3389/fncel.2022.964691.

46. Kole, M.H., and Stuart, G.J. (2008). Is action potential threshold lowest in the axon? Nat Neurosci 11, 1253–1255. 10.1038/nn.2203.

47. Royeck, M., Horstmann, M.T., Remy, S., Reitze, M., Yaari, Y., and Beck, H. (2008). Role of axonal NaV1.6 sodium channels in action potential initiation of CA1 pyramidal neurons. J Neurophysiol 100, 2361–2380. 10.1152/jn.90332.2008.

48. Kole, M.H., Ilschner, S.U., Kampa, B.M., Williams, S.R., Ruben, P.C., and Stuart, G.J. (2008). Action potential generation requires a high sodium channel density in the axon initial segment. Nat Neurosci 11, 178–186. 10.1038/nn2040.

49. Colbert, C.M., Magee, J.C., Hoffman, D.A., and Johnston, D. (1997). Slow recovery from inactivation of Na+ channels underlies the activity-dependent attenuation of dendritic action potentials in hippocampal CA1 pyramidal neurons. J Neurosci 17, 6512–6521. 10.1523/jneurosci.17-17-06512.1997.

50. Park, Y.Y., Johnston, D., and Gray, R. (2013). Slowly inactivating component of Na+ current in peri-somatic region of hippocampal CA1 pyramidal neurons. J Neurophysiol 109, 1378–1390. 10.1152/jn.00435.2012.

51. Toib, A., Lyakhov, V., and Marom, S. (1998). Interaction between duration of activity and time course of recovery from slow inactivation in mammalian brain Na+ channels. J Neurosci 18, 1893–1903. 10.1523/jneurosci.18-05-01893.1998.

52. Miralles, R.M., and Patel, M.K. (2022). It Takes Two to Tango: Channel Interplay Leads to Paradoxical Hyperexcitability in a Loss-of-Function Epilepsy Variant. Epilepsy Curr 22, 69–71. 10.1177/15357597211057966.

53. Lopez-Santiago, L.F., Yuan, Y., Wagnon, J.L., Hull, J.M., Frasier, C.R., O’Malley, H.A., Meisler, M.H., and Isom, L.L. (2017). Neuronal hyperexcitability in a mouse model of SCN8A epileptic encephalopathy. Proc Natl Acad Sci U S A 114, 2383–2388. 10.1073/pnas.1616821114.

54. Sanders, S.J., Campbell, A.J., Cottrell, J.R., Moller, R.S., Wagner, F.F., Auldridge, A.L., Bernier, R.A., Catterall, W.A., Chung, W.K., Empfield, J.R., et al. (2018). Progress in Understanding and Treating SCN2A-Mediated Disorders. Trends Neurosci 41, 442–456. 10.1016/j.tins.2018.03.011.

55. Tran, C.H., Vaiana, M., Nakuci, J., Somarowthu, A., Goff, K.M., Goldstein, N., Murthy, P., Muldoon, S.F., and Goldberg, E.M. (2020). Interneuron Desynchronization Precedes Seizures in a Mouse Model of Dravet Syndrome. J Neurosci 40, 2764–2775. 10.1523/jneurosci.2370-19.2020.

56. Steriade, M., Amzica, F., Neckelmann, D., and Timofeev, I. (1998). Spike-wave complexes and fast components of cortically generated seizures. II. Extra- and intracellular patterns. J Neurophysiol 80, 1456–1479. 10.1152/jn.1998.80.3.1456.

57. Timofeev, I., and Steriade, M. (2004). Neocortical seizures: initiation, development and cessation. Neuroscience 123, 299–336. 10.1016/j.neuroscience.2003.08.051.

58. Lorincz, A., and Nusser, Z. (2008). Cell-type-dependent molecular composition of the axon initial segment. J Neurosci 28, 14329–14340. 10.1523/jneurosci.4833-08.2008.

59. Akin, E.J., Solé, L., Dib-Hajj, S.D., Waxman, S.G., and Tamkun, M.M. (2015). Preferential targeting of Nav1.6 voltage-gated Na+ Channels to the axon initial segment during development. PLoS One 10, e0124397. 10.1371/journal.pone.0124397.

60. Lindy, A.S., Stosser, M.B., Butler, E., Downtain-Pickersgill, C., Shanmugham, A., Retterer, K., Brandt, T., Richard, G., and McKnight, D.A. (2018). Diagnostic outcomes for genetic testing of 70 genes in 8565 patients with epilepsy and neurodevelopmental disorders. Epilepsia 59, 1062–1071. 10.1111/epi.14074.

61. Deng, H., Xiu, X., and Song, Z. (2014). The molecular biology of genetic-based epilepsies. Mol Neurobiol 49, 352–367. 10.1007/s12035-013-8523-6.

62. Agbo, J., Ibrahim, Z.G., Magaji, S.Y., Mutalub, Y.B., Mshelia, P.P., and Mhyha, D.H. (2023). Therapeutic efficacy of voltage-gated sodium channel inhibitors in epilepsy. Acta Epileptologica 5, 16. 10.1186/s42494-023-00127-2.

63. Greenfield, L.J., Jr. (2013). Molecular mechanisms of antiseizure drug activity at GABAA receptors. Seizure 22, 589–600. 10.1016/j.seizure.2013.04.015.

64. Yin, Y.H., Ahmad, N., and Makmor-Bakry, M. (2013). Pathogenesis of epilepsy: challenges in animal models. Iranian journal of basic medical sciences 16, 1119.

65. Encinas, A.C., Watkins, J.C., Longoria, I.A., Johnson, J.P., Jr., and Hammer, M.F. (2020). Variable patterns of mutation density among NaV1.1, NaV1.2 and NaV1.6 point to channel-specific functional differences associated with childhood epilepsy. PLoS One 15, e0238121. 10.1371/journal.pone.0238121.

66. Menezes, L.F.S., Sabiá Júnior, E.F., Tibery, D.V., Carneiro, L.D.A., and Schwartz, E.F. (2020). Epilepsy-Related Voltage-Gated Sodium Channelopathies: A Review. Front Pharmacol 11, 1276. 10.3389/fphar.2020.01276.

67. Strzelczyk, A., and Schubert-Bast, S. (2022). A Practical Guide to the Treatment of Dravet Syndrome with Anti-Seizure Medication. CNS Drugs 36, 217–237. 10.1007/s40263-022-00898-1.

68. Gazina, E.V., Leaw, B.T., Richards, K.L., Wimmer, V.C., Kim, T.H., Aumann, T.D., Featherby, T.J., Churilov, L., Hammond, V.E., Reid, C.A., and Petrou, S. (2015). ‘Neonatal’ Nav1.2 reduces neuronal excitability and affects seizure susceptibility and behaviour. Hum Mol Genet 24, 1457–1468. 10.1093/hmg/ddu562.

69. Boiko, T., Van Wart, A., Caldwell, J.H., Levinson, S.R., Trimmer, J.S., and Matthews, G. (2003). Functional specialization of the axon initial segment by isoform-specific sodium channel targeting. J Neurosci 23, 2306–2313. 10.1523/jneurosci.23-06-02306.2003.

70. Kalume, F., Westenbroek, R.E., Cheah, C.S., Yu, F.H., Oakley, J.C., Scheuer, T., and Catterall, W.A. (2013). Sudden unexpected death in a mouse model of Dravet syndrome. J Clin Invest 123, 1798–1808. 10.1172/jci66220.

71. Oakley, J.C., Cho, A.R., Cheah, C.S., Scheuer, T., and Catterall, W.A. (2013). Synergistic GABA-enhancing therapy against seizures in a mouse model of Dravet syndrome. J Pharmacol Exp Ther 345, 215–224. 10.1124/jpet.113.203331.

72. Bai, Y.F., Zeng, C., Jia, M., and Xiao, B. (2022). Molecular mechanisms of topiramate and its clinical value in epilepsy. Seizure 98, 51–56. 10.1016/j.seizure.2022.03.024.

73. Krook-Magnuson, E., Armstrong, C., Oijala, M., and Soltesz, I. (2013). On-demand optogenetic control of spontaneous seizures in temporal lobe epilepsy. Nature Communications 4, 1376. 10.1038/ncomms2376.

74. Nagaraj, V., Lee, S.T., Krook-Magnuson, E., Soltesz, I., Benquet, P., Irazoqui, P.P., and Netoff, T.I. (2015). Future of seizure prediction and intervention: closing the loop. J Clin Neurophysiol 32, 194–206. 10.1097/wnp.0000000000000139.

75. Ledri, M., Andersson, M., Wickham, J., and Kokaia, M. (2023). Optogenetics for controlling seizure circuits for translational approaches. Neurobiology of Disease 184, 106234.

