## Supplemental Tables and Figures for "Differential roles of Na_V_1.2 and Na_V_1.6 in neocortical pyramidal cell excitability"

**Contents:**

Table 1-4

Key Resources Table

Supplemental Figures 1-6

Supplemental References

**Table 1 – Mutagenic primer sequences for $Na_V$1.2 and $Na_V$1.6**

| Primer name | Sequence (5' to 3') |
|---|---|
| 2A_Y1564S_W1565R_F | ATTCTGT**C**C**C**GGATTAATCTGGTGTTTATTGTTCT |
| 2A_Y1564S_W1565R_R | AGATTAATCC**G**G**G**ACAGAATGTTTGTCATTTCTTGA |
| 8A_Y1555S_W1556R_F | CATCCTCT**C**C**C**GGATTAACCTGGTGTTTGTT |
| 8A_Y1555S_W1556R_R | GGTTAATCC**G**G**G**AGAGGATGTTCTCCATCTG |

**Table 2 – Voltage dependence of activation and inactivation for wildtype and mutant $Na_V$ isoforms, related to Figure 1.**

| $Na_V$ channel | V ½ of activation (mV) | Slope factor | V ½ of inactivation (mV) | Slope factor |
|---|---|---|---|---|
| $Scn2a^{YW/YW}$ or Wildtype $Na_V$1.2 (n=6) | -20.1 ± 2.6 | 6.6 ± 0.5 | -68.3 ± 1.8 | 6.9 ± 0.3 |
| $Scn2a^{SR/SR}$ or Mutant $Na_V$1.2 (n=7) | -18.7 ± 2.1 | 6.4 ± 0.6 | -68.1 ± 0.8 | 7.1 ± 0.5 |
| $Scn8a^{YW/YW}$ or Wildtype $Na_V$1.6 (n=8) | -23.9 ± 2.3 | 6.7 ± 0.4 | -74.6 ± 3.3 | 6.9 ± 0.5 |
| $Scn8a^{SR/SR}$ or Mutant $Na_V$1.6 (n=6) | -26.1 ± 3.2 | 6.5 ± 0.4 | -71.9 ± 1.2 | 7.0 ± 0.3 |

**Table 3 – Neuronal AP firing properties at baseline, related to Figures 3 & 4**

| $Na_V$ channel | Threshold (mV) | Peak dV/dt (V/s) | Spike # | Last AHP |
|---|---|---|---|---|
| No drug control (n=12) | -42.53 ± 0.61 | 555.82 ± 17.57 | 5.83 ± 0.41 | -49.47 ± 0.43 |
| Wildtype (n=11) | -42.41 ± 0.83 | 554.42 ± 14. 5 | 5.36 ± 0.36 | -50.04 ± 0.71 |
| $8a/2a^{SR/SR}$ (n=12) | -44.87 ± 0.47 | 545.11 ± 12.89 | 5.09 ± 0.16 | -51.39 ± 0.5 |
| $Scn2a^{SR/SR}$ (n=12) | -42.3 ± 0.76 | 520.05 ± 13.51 | 6.09 ± 0.32 | -49.31 ± 0.84 |
| $Scn8a^{SR/SR}$ (n=12) | -43.87 ± 0.65 | 517.36 ± 21.83 | 5.18 ± 0.55 | -50.46 ± 0.74 |

**Table 4 – AP firing properties of first spike in 10 sec AP train, related to Figures 5 & 6**

| $Na_V$ channel | Threshold (mV) | Peak dV/dt (V/s) |
|---|---|---|
| Wildtype, no drug (n=12) | -43.1 ± 0.4 | 549.1 ± 8.8 |
| Wildtype + GNE-4076 (n=12) | -41.7 ± 0.7 | 541.0 ± 12.5 |
| $8a/2a^{SR/SR}$ + GNE-4076 (n=12) | -43.7 ± 0.4 | 561.3 ± 14.5 |
| $Scn2a^{SR/SR}$ + GNE-4076 (n=10) | -40.7 ± 0.4 | 500.7 ± 12.5 |
| $Scn8a^{SR/SR}$ + GNE-4076 (n=12) | -43.3 ± 0.5 | 542.9 ± 11.6 |

**Key Resources Table**

| REAGENT or RESOURCE | SOURCE | IDENTIFIER |
|---|---|---|
| **Chemicals, peptides and recombinant proteins** | | |
| GNE-4076 or Compound 5[1] | Merck & Co., Inc. | Roecker et al.[1]; PMID: 28389149 |
| **Experimental Models: Organisms/Strains** | | |
| Mouse: C57B6J | The Jackson Laboratory | RRID: IMSR_JAX:000664 |
| Mouse: $Na_V$1.6 YW→SR KI | Genentech | Deng et al.[2]; PMID: 37352856 |
| Mouse: $Na_V$1.2 YW→SR KI | Genentech; This manuscript | N/A |
| Mouse: Dual $Na_V$1.2/1.6 KI YW→SR KI | Genentech; This manuscript | N/A |
| **Oligonucleotides** | | |
| 2A_Y1564S_W1565R F: ATTCTGTCCCGGATTAATCTG GTGTTTATTGTTCT | This manuscript | N/A |
| 2A_Y1564S_W1565R R: AGATTAATCCGGGACAGAATG TTTGTCATTTCTTGA | This manuscript | N/A |
| 8A_Y1555S_W1556R F: CATCCTCTCCCGGATTAACCT GGTGTTTGTT | This manuscript | N/A |
| 8A_Y1555S_W1556R R: GGTTAATCCGGGAGAGGATGT TCTCCATCTG | This manuscript | N/A |
| **Recombinant DNA** | | |
| pIR-CMV-SCN2A-Variant-1-IRES-mScarlet | Addgene | RRID: Addgene_162279 |
| pcDNA4TO-SCN8A-Variant-3-IRES-mScarlet | Addgene | RRID: Addgene_209411 |
| **Software** | | |
| IGOR Pro v6.3 & v9 (Wavemetrics) | https://www.wavemetrics.com/ | RRID: SCR_000325 |
| Prism 10 (GraphPad) | https://www.graphpad.com/scientific-software/prism/ | RRID: SCR_002798 |
| NEURON Compartmental Models (modelDB) | https://www.neuron.yale.edu/neuron/ | RRID: SCR_007271 & SCR_003105 |
| Python | https://www.python.org | RRID: SCR_008394 |
| Benchling | https://www.benchling.com | RRID: SCR_013955 |

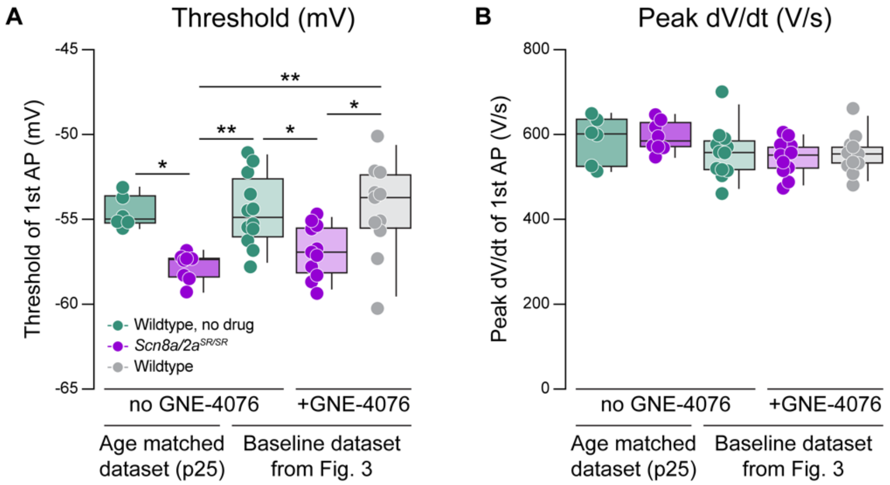


**Supplemental Figure 1: Dual knock-in mutants (Scn8a/2aSR/SR) hyperpolarizes AP threshold without impairing peak dV/dt, related to Figure 1**

(A) Summary data for threshold (mV) with or without bath application of GNE-4076. Data was collected from age matched groups (without GNE-4076) or analyzed from baseline data presented in Figure 3. Box plots show median and 90% tails. Circles represent individual cells (wildtype no drug at p25, n=6; 8a/2aSR/SR no drug at p25, n=8; wildtype no drug from Fig. 3, n=12; 8a/2aSR/SR + GNE-4076 from Fig. 3, n=11; wildtype + GNE-4076 from Fig. 3, n=11). One-way ANOVA, Holm-Šídák multiple comparisons test.*p<0.05, *p<0.05, **p<0.01.

(B) Summary data for peak dV/dt (V/s) with or without bath application of GNE-4076. Data was collected from age matched groups (without GNE-4076) or analyzed from baseline data presented in Figure 3. Box plots show median and 90% tails. Circles represent individual cells One-way ANOVA, Holm-Šídák multiple comparisons test. n.s.

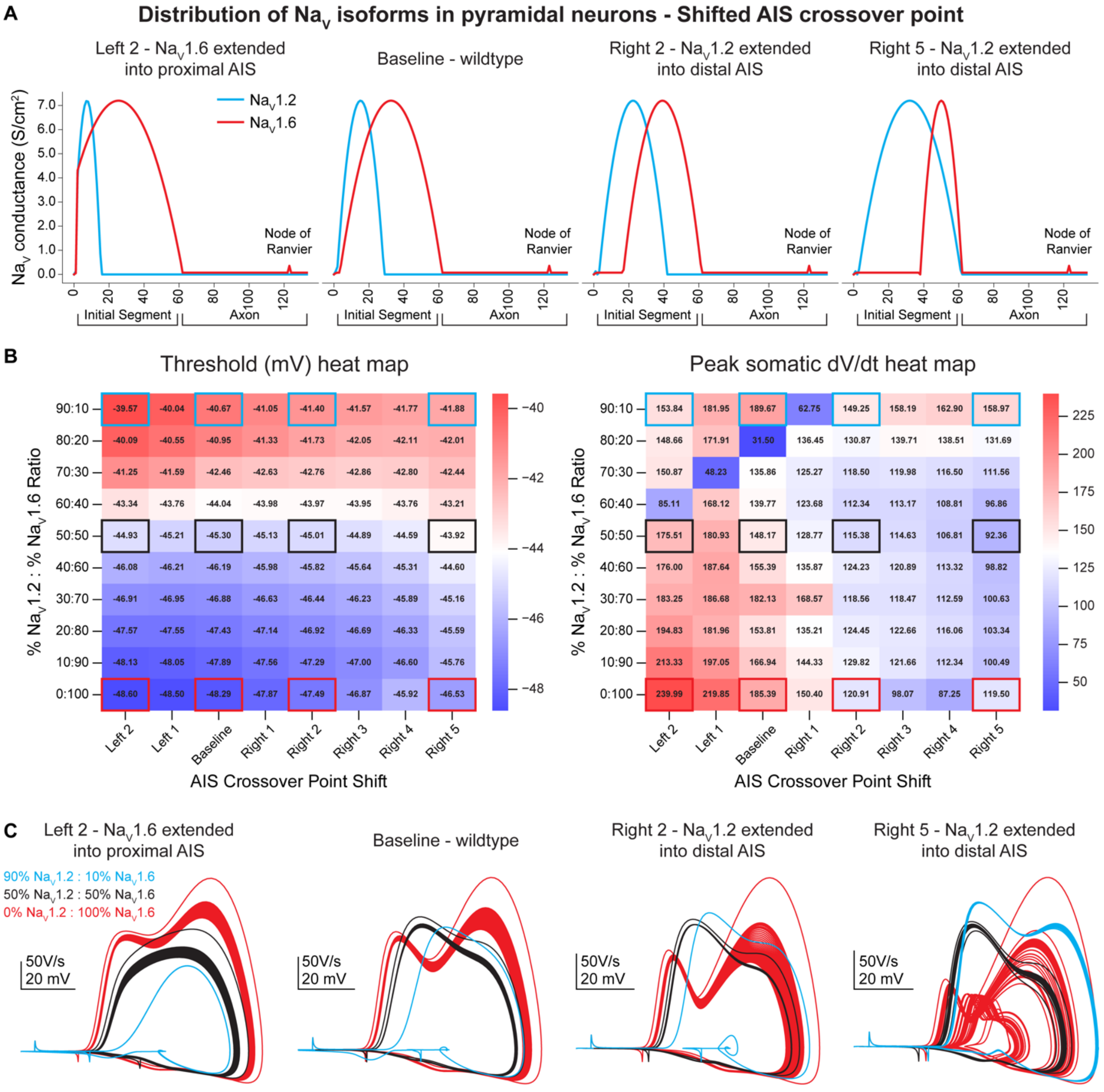


**Supplemental Figure 2: Compartmental model sensitivity analysis, related to Figure 2**

(A) To assess model sensitivity, NaV1.6 and NaV1.2 distribution is shifted throughout the axon initial segment (AIS). The AIS crossover point is adjusted from baseline with NaV1.6 extended into the proximal AIS (left 2) or NaV1.2 extended into the distal AIS (right 2 and 5).

(B) Heat maps for AP threshold and peak somatic dV/dt at different AIS crossover points. NaV1.2 and NaV1.6 ratios are also varied along the y-axis.

(C) Example phase plane traces at different AIS crossover positions. Each AIS crossover position shows individual plots based on the different NaV ratios. Increased NaV1.2 density consistently depolarized AP threshold while dV/dt peaks are randomly altered at the different density ratios. Scale bars are unique to each AIS crossover position.

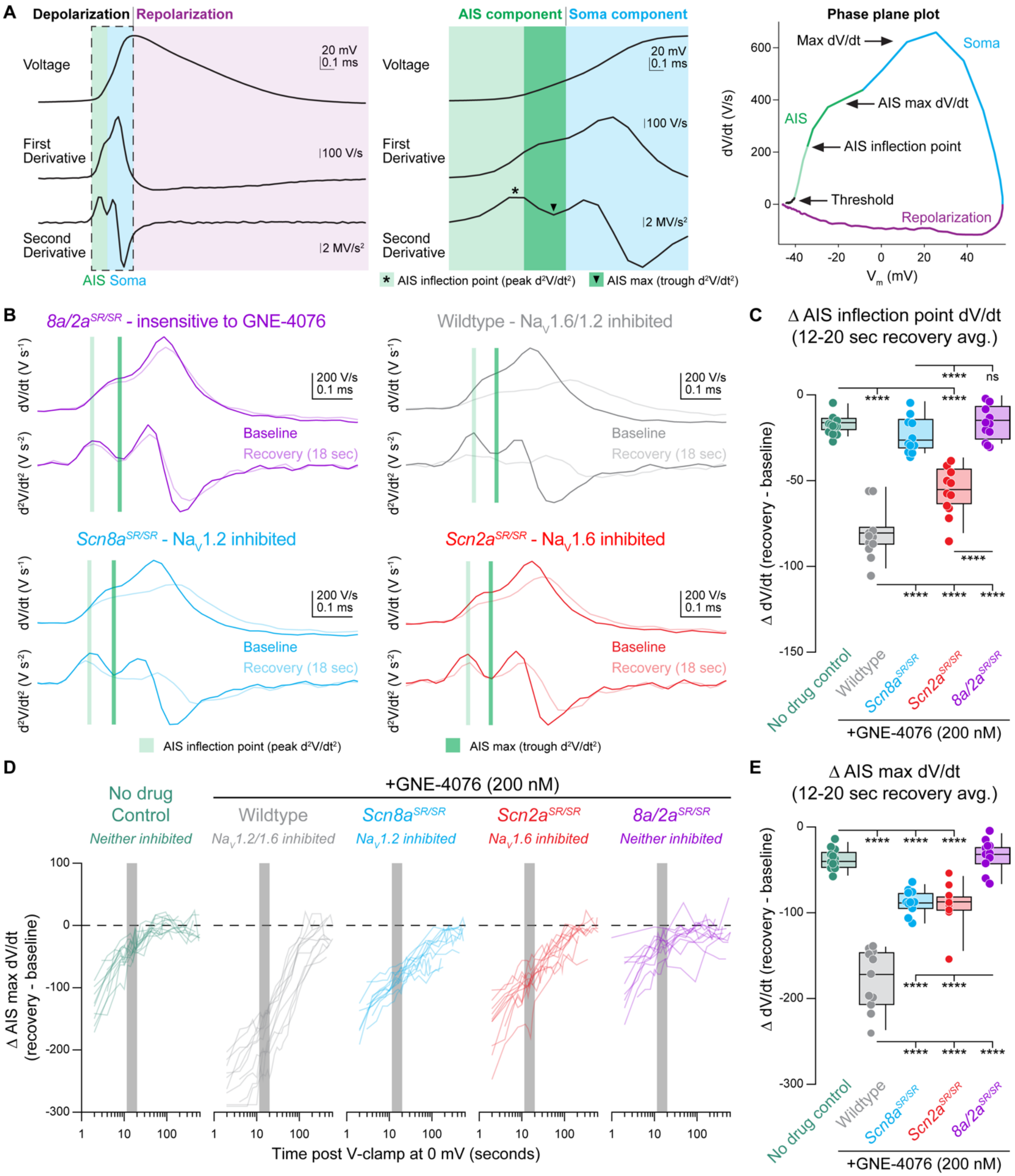


**Supplemental Figure 3: Depolarization across the AIS compartment is mediated by distinct NaV isoforms, related to Figure 3**

(A) Schematic depicting different phases of an action potential (AP), including AP velocity (1st derivative; dV/dt or V/s) and acceleration (2nd derivative; d2V/dt2 or V/s2). APs can be divided based on depolarization or repolarization phases, which largely depends on opening of various voltage-gated ion channels. During cellular depolarization, two components exist that largely account for voltage change detected at either the AIS or soma, which are easily discerned by two humps present on a phase plane plot. The AIS component can further be separated into the AIS max or inflection point defined as the trough or peak of the second derivative and likely represents voltage change at distinct AIS regions like the proximal or distal AIS, respectively.

(B) Example traces of 1st and 2nd derivatives for all genotypes at baseline or 18 sec post recovery. AIS inflection point (light green) is defined as corresponding dV/dt when acceleration (d2V/dt2) peaks during the AIS component. AIS max point (dark green) is defined as corresponding dV/dt when acceleration plateaus (visualized as a trough of d2V/dt2) during AIS component.

(C) Summary data for D AIS inflection point (dV/dt) at 12-20 sec post GNE-4076 onboarding (represented as light green bar in (B)). Box plots show median and 90% tails. Circles represent individual cells (wildtype no drug, n=12; wildtype + GNE-4076, n=11; Scn2aSR/SR + GNE-4076, n=11; Scn8aSR/SR + GNE-4076, n=11; 8a/2aSR/SR + GNE-4076, n=11). One-way ANOVA, Holm-Šídák multiple comparisons test. ****p<0.0001.

(D) Recovery of AIS max velocity (dV/dt) represented as a delta value for individual cells plotted against time post GNE-4076 onboarding (log-scale). For D dV/dt, baseline value is subtracted from individual timepoints throughout the recovery phase (D V/s= recovery timepoint - baseline). Gray shaded bar represents recovery between 12-20 sec.

(E) Summary data for D AIS max velocity (dV/dt) at 12-20 sec post GNE-4076 onboarding (represented as gray bar in (D) or dark green bar in (B)). Box plots show median and 90% tails. Circles represent individual cells. One-way ANOVA, Holm-Šídák multiple comparisons test. ****p<0.0001.

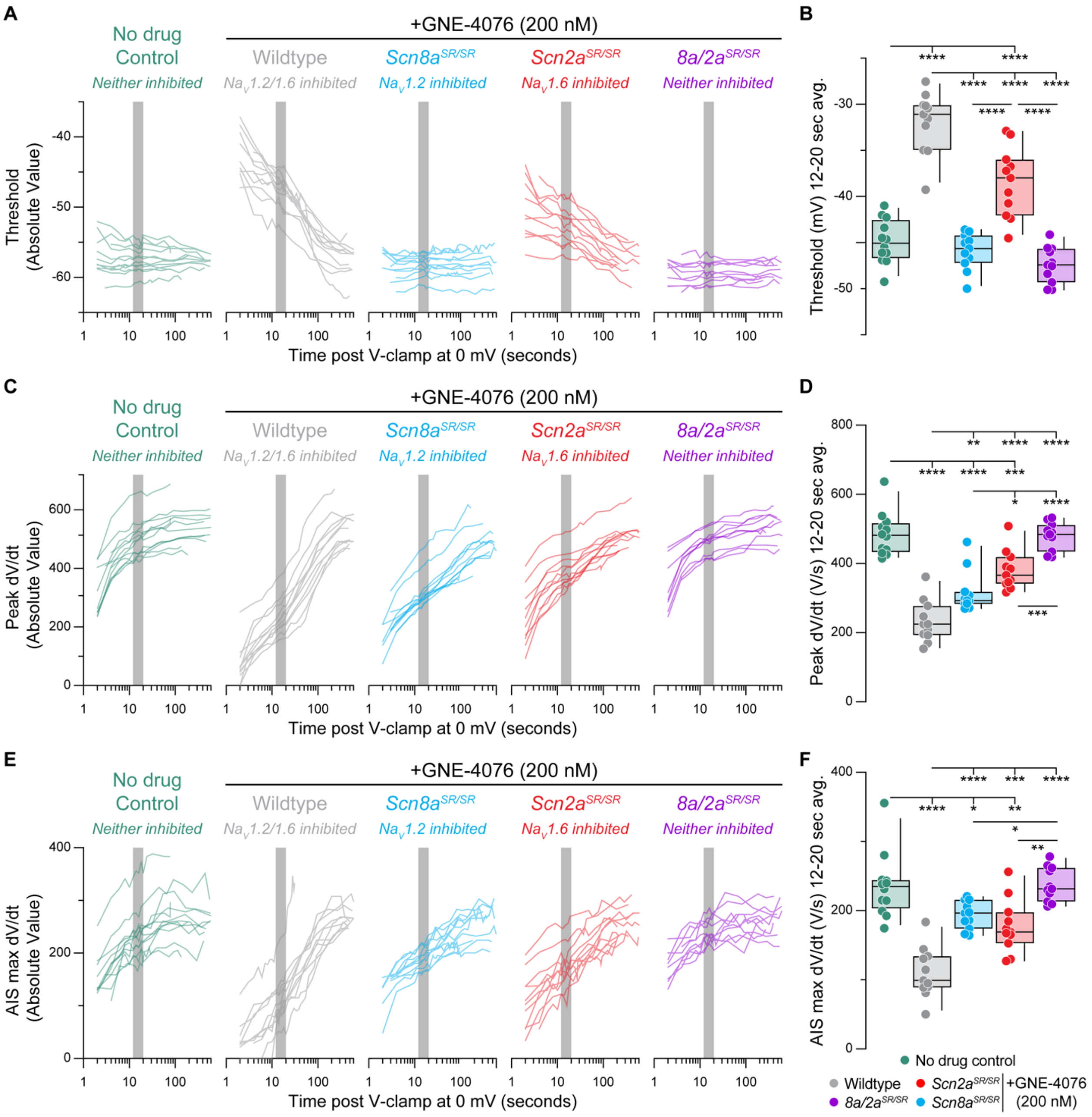


**Supplemental Figure 4: Recovery of AP firing properties represented as absolute values, related to Figure 3 and Supplemental Figure 3**

(A) Recovery of AP threshold (Vm) represented as absolute values for individual cells (wildtype no drug, n=12; wildtype + GNE-4076, n=11; Scn2aSR/SR + GNE-4076, n=11; Scn8aSR/SR + GNE-4076, n=11; 8a/2aSR/SR + GNE-4076, n=11) plotted against time post GNE-4076 onboarding (log-scale). Colors are matched to conditions represented in Fig. 3B. Gray shaded bar represents recovery between 12-20 sec.

(B) Summary data for Vm at 12-20 sec post GNE-4076 onboarding (time period represented as gray bar in (A)). Box plots show median and 90% tails. Circles represent individual cells. One-way ANOVA, Holm-Šídák multiple comparisons test. *p<0.05, **p<0.01, ***p<0.001, ****p<0.0001.

(C) Recovery of AP peak velocity (dV/dt) represented as absolute values for individual cells plotted against time post GNE-4076 onboarding (log-scale). Colors are matched to conditions represented in Fig. 3B. Gray shaded bar represents recovery between 12-20 sec.

(D) Summary data for peak dV/dt at 12-20 sec post GNE-4076 onboarding (time period represented as gray bar in (C)). Box plots show median and 90% tails. Circles represent individual cells. One-way ANOVA, Holm-Šídák multiple comparisons test. *$p<0.05$, **$p<0.01$, ***$p<0.001$, ****$p<0.0001$.

(E) Recovery of AIS max velocity (dV/dt) represented as absolute values for individual cells plotted against time post GNE-4076 onboarding (log-scale). Colors are matched to conditions represented in Fig. 3B. Gray shaded bar represents recovery between 12-20 sec.

(F) Summary data for AIS max velocity (dV/dt) at 12-20 sec post GNE-4076 onboarding (time period represented as gray bar in (E)). Box plots show median and 90% tails. Circles represent individual cells. One-way ANOVA, Holm-Šídák multiple comparisons test. *$p<0.05$, **$p<0.01$, ***$p<0.001$, ****$p<0.0001$.

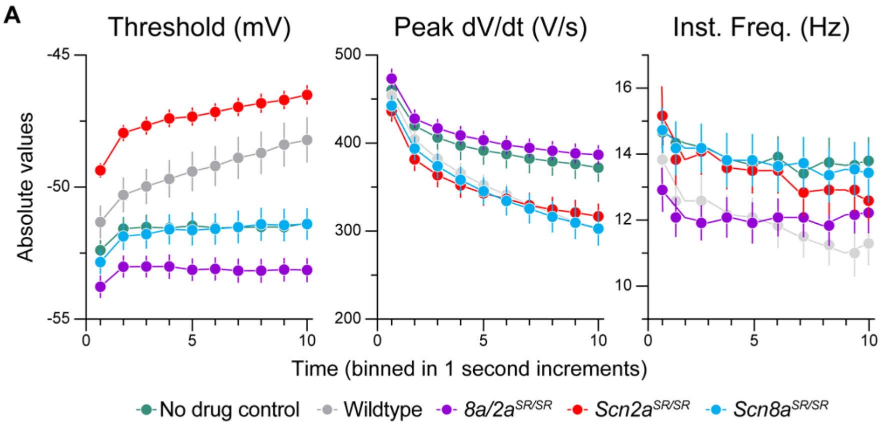


**Supplemental Figure 5: Absolute values for activity-dependent onboarding of GNE-4076, related to Figure 5 and 6**

(A) Absolute values for threshold (mV), peak dV/dt (V/s) and instantaneous firing frequency (Hz) binned in 1 sec increments. Circles represent mean absolute value ± SEM (wildtype no drug, n=12; wildtype + GNE-4076, n=12; 8a/2aSR/SR + GNE-4076, n=12; Scn2aSR/SR + GNE-4076, n=10; Scn8aSR/SR + GNE-4076, n=12). Two-way ANOVA, Holm-Šídák multiple comparisons test.

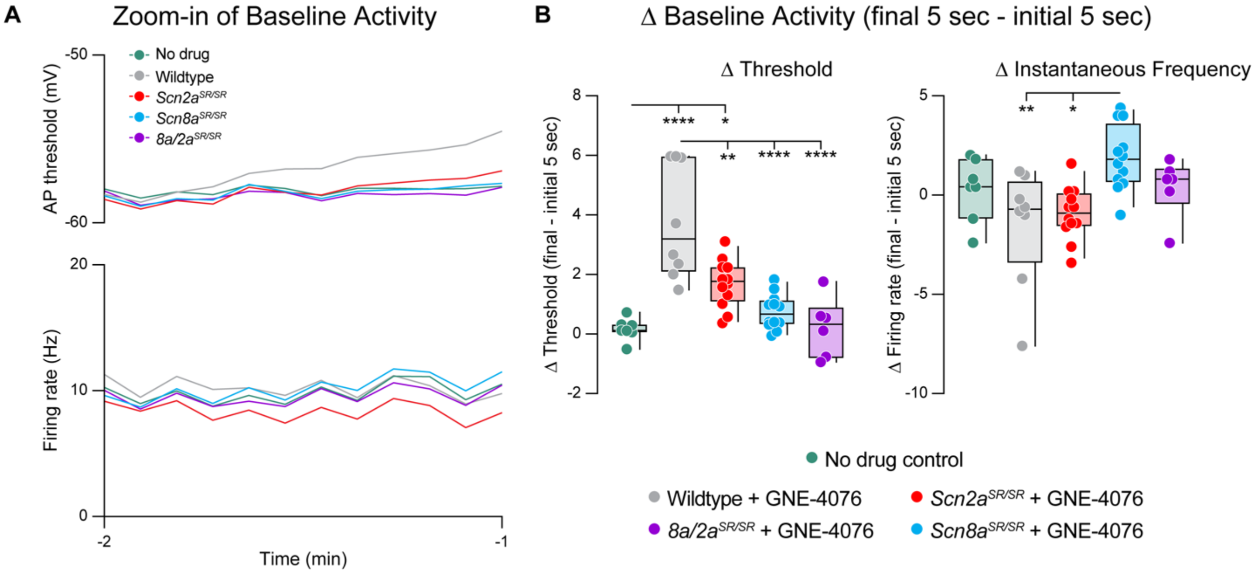


**Supplemental Figure 6: GNE-4076 onboarding during baseline post-synaptic potential (PSP) activity, related to Figure 7**

(A) Changes to baseline activity prior to seizure-like activity (Zoom-in of Fig. 7C). Absolute values for both threshold (mV) and instantaneous firing frequency (Hz) are binned in 5 sec increments for all genotypes in Fig. 7B.

(B) Summary of threshold or instantaneous frequency data for the final 5 sec of baseline activity in (A). Delta values are normalized to the initial 5 sec of baseline activity (final - initial 5 sec). Box plots show median and 90% tails. Circles represent individual cells (wildtype no drug, n=7; wildtype + GNE-4076, n=8; Scn2aSR/SR + GNE-4076, n=12; Scn8aSR/SR + GNE-4076, n=12; 8a/2aSR/SR + GNE-4076, n=6). One-way ANOVA, Holm-Šídák multiple comparisons test. *p<0.05, **p<0.01, ****p<0.0001.

**Supplemental References:**


1. Roecker, A.J., Egbertson, M., Jones, K.L.G., Gomez, R., Kraus, R.L., Li, Y., Koser, A.J., Urban, M.O., Klein, R., Clements, M., et al. (2017). Discovery of selective, orally bioavailable, N-linked arylsulfonamide Na(v)1.7 inhibitors with pain efficacy in mice. Bioorg Med Chem Lett *27*, 2087-2093. 10.1016/j.bmcl.2017.03.085.
2. Deng, L., Dourado, M., Reese, R.M., Huang, K., Shields, S.D., Stark, K.L., Maksymetz, J., Lin, H., Kaminker, J.S., Jung, M., et al. (2023). Nav1.7 is essential for nociceptor action potentials in the mouse in a manner independent of endogenous opioids. Neuron *111*, 2642-2659.e2613. 10.1016/j.neuron.2023.05.024.
